# Towards Environmental Control of Microbiomes

**DOI:** 10.1101/2022.11.04.515211

**Authors:** Will Sharpless, Kyle Sander, Fangchao Song, Jennifer Kuehl, Adam Paul Arkin

## Abstract

Microbial communities have consequential effects on health and the environment yet remain uncontrollable due to their complex dynamics. Ecological modeling offers a platform to overcome their nonlinear and interconnected nature but traditionally does not account for context-dependence. Here, we extend the generalized Lotka-Volterra (gLV) model to accommodate a varying environment by identifying how environmental changes alter species growth rates and interactions in a manner that predicts full community trajectories across environmental gradients. We identify key environment-varying interactions within a synthetic community derived from the Oryzae sativa rhizosphere, and demonstrate how variations in the environment change fixed point compositions and rates of convergence. With our model, we simulate how precise perturbations of the environment can offer improvements in an optimal control problem of driving a community to a target composition. We show that environmental perturbation can minimize the total species input (direct species perturbation) and greatly expand the set of initial states from which a desired target can be reached despite stochasticity. This work demonstrates that a formal perspective on environmental influence of community dynamics is valuable for not only understanding seasonal changes or anthropogenic manipulations, but is critical for improving control of the microbiome.

## Introduction

Microbiome states have been correlated with health and disease across biological kingdoms. Specific diversities and abundances of microbes have specific metabolic potentials that influence obesity, immunity, and cognition in humans [8,9,10] as well as pathogen resistance, drought resilience and yield in agriculture [11]. There is a desire to understand the mechanisms by which the microbiome responds to and affects its environments. Models have arisen which are used to infer and explain observed microbiome population dynamics and activity. While these range from complex multi-organism metabolic network models [12, 13] to highly abstracted trait-based models [14], the generalized Lotka-Volterra (gLV) remains one of the more powerful and popular frameworks for understanding how interactions among microbiome members leads to observed dynamics, with predictive successes in 3 to 25 member microbial communities [15,16,17]. This model takes the form,

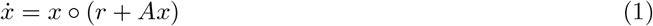

where *x* ∈ ℝ^*n*^ is the vector of species population sizes (OD/mL), *r* ∈ ℝ^*n*^ is the vector of innate growth rates, and *A* ∈ ℝ^*n×n*^ is the matrix of all pairwise interactions (with *α*_*ij*_ being the effect of *x*_*j*_ on *x*_*i*_’s growth and *α*_*ii*_ being the effect of *x*_*i*_ on itself). Note, ∘ is used to represent element-wise multiplication. Given the accuracy of these models [15,16,17], it seems reasonable to ask if they could be used to predict necessary interventions to achieve a desired microbiome state, for example, to recover from a state inducing disease and restore a state corresponding to a beneficial phenotype.

Theorists have proposed a control affine model for manipulating gLV systems,

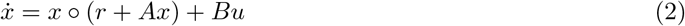

based on inputs *u* ∈ ℝ^*m*^ of probiotics or antibiotics with respective sensitivities *B* ∈ ℝ^*n×m*^ [18]. Notably, Angulo *et al*. 2019 drew from the theory of structural controllability [19] to write a graph-based algorithm for identifying a minimal set of ‘driver species’ (inputs) to make (2) fully controllable, and then demonstrated its validity *in silico* with a linearized Model Predictive Controller (MPC) with a high success rate. However, this framework operates in a fixed context by assuming that innate growth rates *r*, interactions *A*, and input sensitivities *B* remain constant throughout control programs. In this work, we expand this fixed model to demonstrate how environmentally-varying parameters are not only accurate but improve controller performance.

Research suggests that fixed growth rates and interactions is a tenuous assumption if the environment is subject to fluctuation [20, 21, 22], and here we consider this complexity to interrogate whether it might improve understanding and frameworks for microbiome control. For example, bacteria can sense ecological competition via nutrient limitation and respond with antibiotic production and other antagonistic measures [21, 22], and thus, the addition of nutrients including prebiotic molecules like carbohydrates should attenuate or neutralize negative interspecies interactions. As the environment becomes richer, and antagonistic pressures theoretically subside, we might expect a reduction in interaction density in the community and an increase network partitioning, that alters the stable composition and requires a larger driver species set.

While this extension may seem complex, regarding these interactions as variable may be valuable for driving gLV systems for the same reason. Different ecological states may yield more amenable ecological dynamics for control offering faster convergence, alternative driver species, or fewer required input interventions. These kinds of benefits could simplify modulation programs in medical microbiome therapies as well as suggest fertilizer schedules to prime crop rotations.

We test these ideas with a community composed of O. sativa rhizosphere isolates with V4-V5 16S regions most closely related to *Rhizobium pusense* (RP), *Shinella kummerowiae* (SK), *Microbacterium phyllosphaerae* (MP), *Pseudomonas koreensis* (PK), *Bacillus megaterium* (BM), *Pantoea agglomerans* (PA), and *Flavobacterium ginsengiterrae* (FG). We note that two of the isolates, closest to SK and FG, are distantly related to these nearest known organisms and thus we call them iSK and iFG to underscore their lack of relation (Table S1), otherwise all isolates have greater than 98% sequence identity to their nearest relative are called by the same abbreviation. Isolates were chosen to include members with genetic similarity to Plant-Growth-Promoting or antagonistic species, members which are frequently abundant in the rhizosphere, as well as members with a range of growth rates (Figure 1, Table S1). We included the distantly known isolates iSK and iFG to simulate the realistic dilemma of a bioengineer designing a microbial community with local microbes at hand.

**Figure 1:**
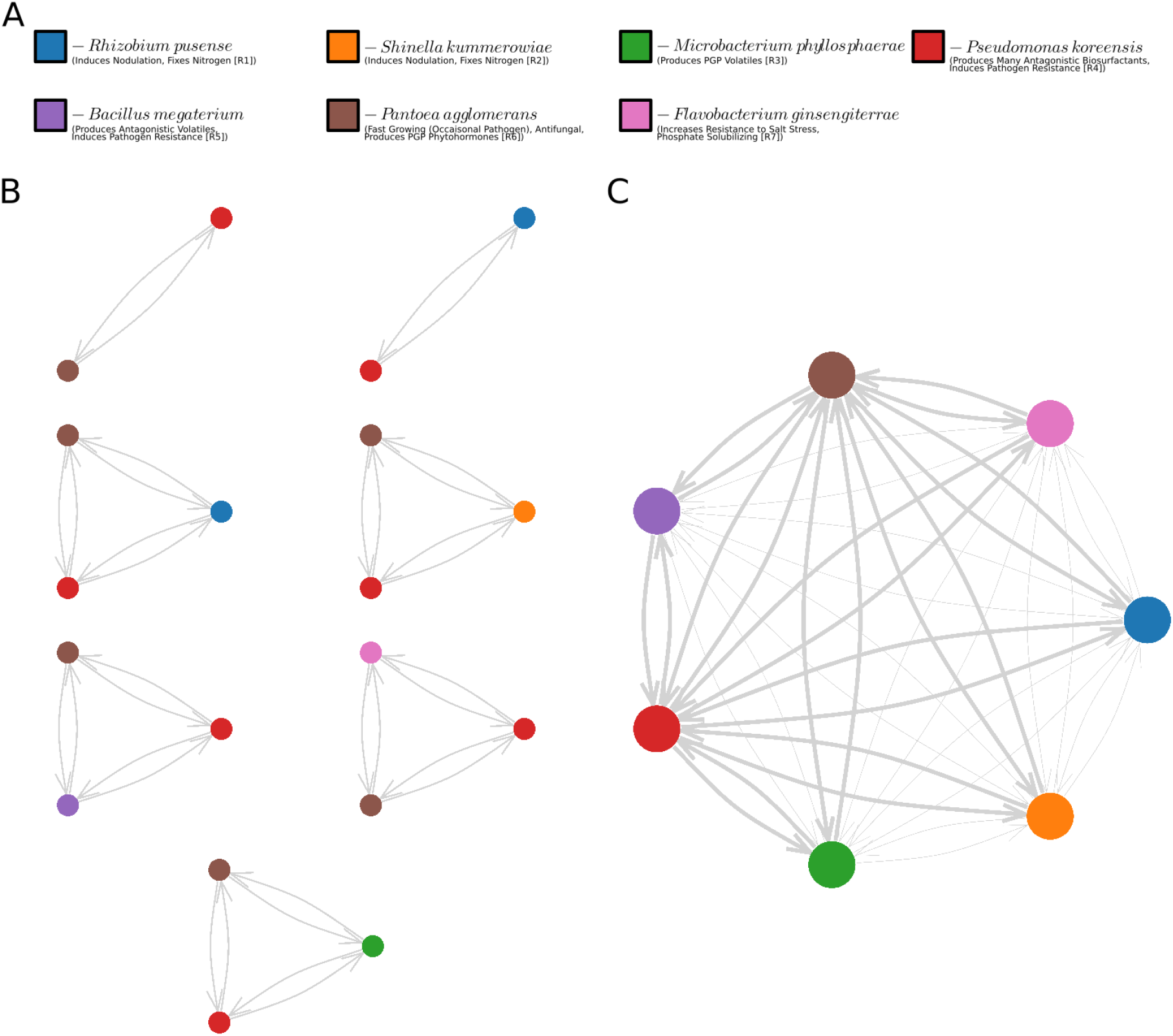
**A**. Legend of isolate-related species (colors consistent in all figures, see Table S1 for strain details) and relevance to rhizosphere microbiome design [1,2,3,4,5,6,7]. Note, the isolates most closely related to SK and FG respectively are distant and might not possess the known abilities. **B**. The training set of subcommunities cultured to parameterize full community model. **C**. Resulting gLV network approximation (65% of all parameters) used to model the full community culture growth. Bold edges represent parameterized interactions and thin edges represent unmeasured interactions modeled with coefficient zero.

Cultures were grown in a liquid phase and passaged to mimic the continuous recirculating environment of a hydroponic rhizosphere where nutrients are periodically replaced. Subsets of the organisms were grown in isogenic, pairwise and triwise cultures to parameterize a rigorous model similar to that demonstrated by Venturelli *et al*. 2019. However, we limited subcommunity cultures for parameterization to all single organisms, pairs between the most abundant organisms, and triplicates of all other organisms with the two most abundant members, to minimize the amount of required training data for the model. We show that this amount and type of training data proved sufficient to accurately predict microbiome trajectories in full community cultures along the environmental gradients of temperature and glucose in static and switched contexts (Figure 3).

We chose to culture the community along two environmental gradients: initial glucose concentration and temperature. It is well known that plants under duress already utilize population-directed environmental control by exuding certain carbohydrates to increase the abundance of PGP species [4]. Alternatively, temperature is an easily controlled variable from an engineering standpoint with global effects on chemical kinetics. We hypothesized that a parameter like glucose might induce discrete changes in physiological states through mechanisms such as catabolite repreression, resulting in hybrid system dynamics [23, 24], while temperature would affect kinetic reaction rates universally and gradually, inciting continuous shifts in the dynamics. In either case, changes in the environment are assumed to impact growth rates *r* and interactions *A* captured by the gLV model. Ultimately, the experiments demonstrate how the given microbial community dynamics vary on environmental gradients and stabilize to various diversities, suggesting the significance and usefulness of an environment based gLV model with accurately identified context-dependent parameters.

We use the validated model to explore *in silico* how this understanding could improve microbiome control programs (ie. therapies for altering composition) in both continuous and hybrid environment-varying contexts. To theoretically drive the studied synthetic consortia, we use Model Predictive Control (MPC) [25] as well as Hamilton-Jacobi-Isaacs (HJI) reachability analysis [26] on the following stochastic gLV formulation,

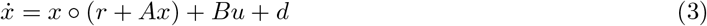

with bounded random disturbance *d* ∈ ℝ^*n*^. Both control programs are evaluated with and without the autonomous decision to vary growth rates *r* and interactions *A*. While limited to simulation, these last analyses underscore the influence of the environment in the design of microbial therapies. Scheduled manipulations of the environment can result in changes in growth rates and interactions that reduce the necessary species input in a therapeutic program as well as increase the radius of treatable states making microbiome control more efficient and effective.

Here we set out to examine the impact of changeable environmental parameters on the dynamics and control of microbial communities using a 7-member consortium of bacteria isolated from the rhizosphere of *Oryza sativa* (rice). To this end, we first parameterize the gLV dynamics of the consortium with isogenic and multi-species subcommunities (Figure 1) cultured along gradients of initial glucose concentration and temperature. We validate the model by testing the fit with the full community cultured in static and switched environments. We analyze the variations in the validated model to identify key environment-varying interactions and trends in growth rates and interactions over environmental gradients (Figure 2). We also consider how the environment changes the dynamics of the system through the change of the full community fixed point and the rate of convergence towards it. Finally, with the parameterized models, we simulate autonomous control of the microbiome to demonstrate how environment-variation can reduce the total species input needed and expand the set of controllable initial states.

**Figure 2:**
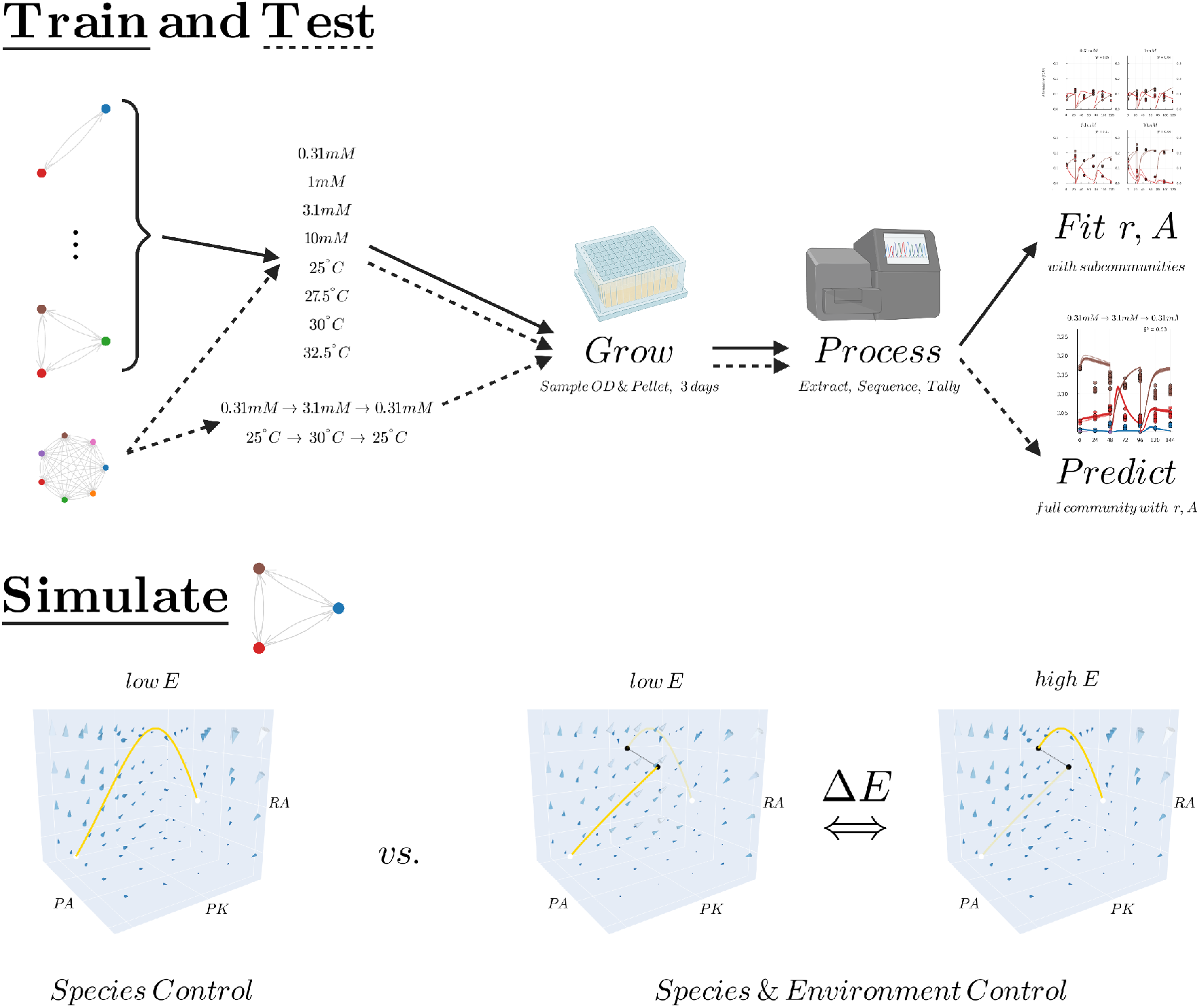
Process Graphic. (Top) First, two environment-varying gLV models were trained by culturing the subcommunities in Fig 1 in various glucose concentrations and temperatures for three days, sampling culture for sequencing and OD to process, sequence and combine into absolute-abundance, training data. Next, these models were tested on data identically garnered from the full community cultured in the same conditions and two additional switched conditions. (Bottom) With the environment-varying models of the principle subcommunity RA-PK-PA, two autonomous, optimal controllers were tested with and without the ability to control the environment (in addition to the ability to actuate species) to interrogate the utility of an environment-varying model.

## Results & Discussion

### Precise Change of the Environment Produces Predictable Change in Community Dynamics

The best-fit model (Table S2) predicts the full community trajectories with 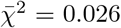 & 0.042 for glucose and temperature gradients respectively (Figure 3). The fits were accomplished with data for only half the cultures compared to the number required for a full pairwise parameterization, but we note that the full community transitions to a subcommunity set which was included in the training data (Methods). The microbiome at hand converges to states where isolates closely related to PA, PK and RA occupy *≥* 99% of the total abundance while members iSK, MP, BM transiently appear at 0.1 to 1% and iFG appears at *<* 0.1%. At higher glucose concentration and temperature, the abundances of RA and PK drop while PA further out-competes PK and RA leaving a nearly isogenic microbiome. In the glucose variation model, Sobol sensitivity analysis of the best fit parameters supports that the interactions between PA, PK and RA are the basis of the community trajectory and outcome (Figures S14). The Sobol analysis of the temperature variation model yields a more uniform distribution of sensitivity despite also producing high fidelity fits (Figure S15), perhaps because of the flexibility of the gLV model. We note that, unlike previous methods [15], the freedom of our training regime to simultaneously weigh isogenic and pairwise data resulted in mediocre fits to the isogenic growth cultures in the training data for the Glucose system and poor fits for the Temperature system (Figure S13), yet strong generalization to the test data in both cases (Figure 3). Although, the good generalization could be due to the full community convergence to a subcommunity within the training set (RA-PK-PA), this could also be due to the well known difference in ecological behavior between isolated organisms and those that sense competition [21]. Regardless, the accuracy of the parameterized glucose model suggests at least that differences in population dynamics across some environmental gradients is explained by changes in ecological interactions between species and with themselves.

**Figure 3:**
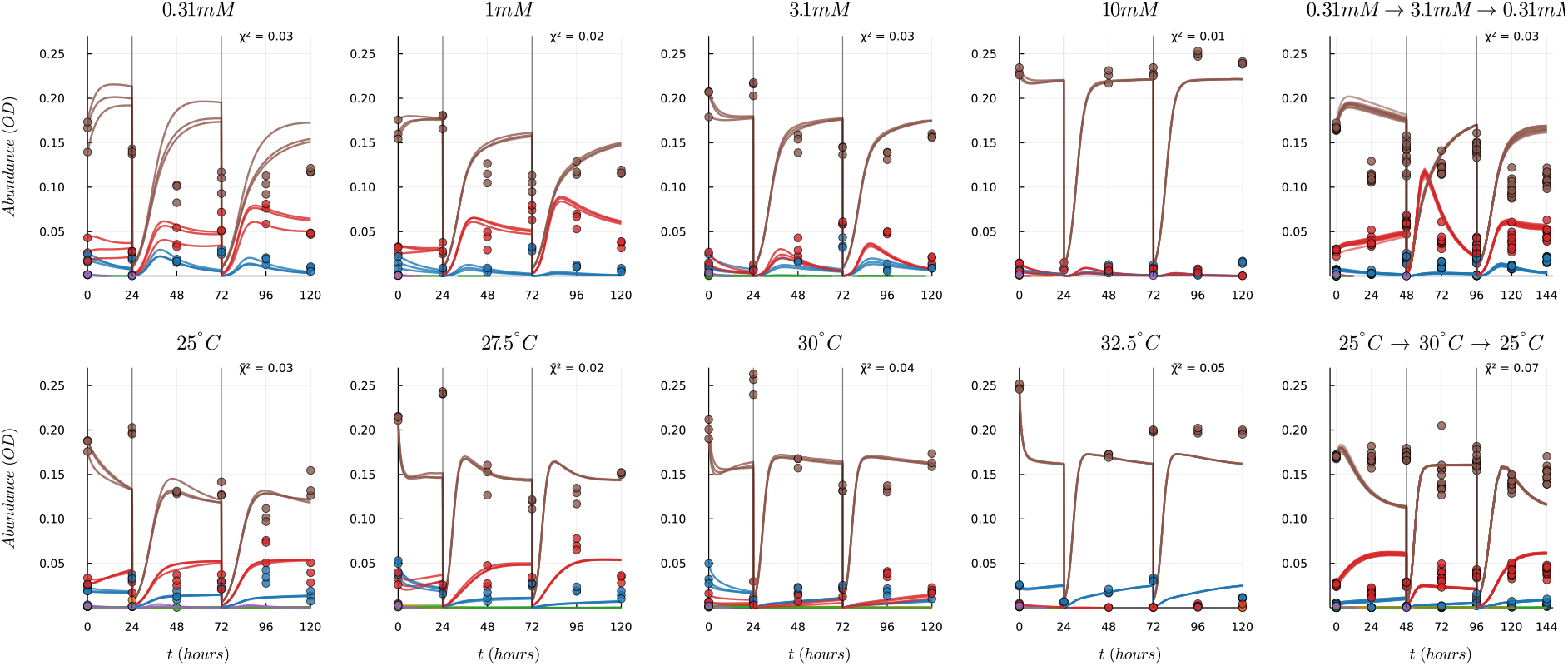
Model Validation with Best Fit. Data (circles) vs. simulations (lines) of full-community systems in static or switched environments. Mean goodness-of-fit 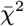 for all species and replicates listed in the upper right corner.

Of the pairwise data, variation in interspecies dynamics over environmental gradients is most noticeable between members PK and PA (Figure 11). At lower glucose and temperature, the two species balance one another and, across replicates, the relative abundance switches back and forth around 1:1 over the 24 hour time-points. However, as glucose concentration or temperature increases, the relative abundance diverges and PA out-competes PK by 144 hours, despite both having improved growth in the corresponding isogenic settings (Figure 11). While not true for all species, the outcome of these two principal members matches the ranking of their isogenic fixed-point abundances in all environment settings. The best-fit model captures this with *−r*_*PA*_*/α*_*PAPA*_ *> −r*_*PK*_*/α*_*PKPK*_ and suggests that strong negative pressures *α*_*PA−PK*_ and *α*_*PK−PA*_ allow one another foothold. The observed trends in interactions reiterate the previously postulated principle that the best competitive advantage is rapid and massive growth in rich circumstances and negative interactions stabilize a diverse community [27]. Ultimately, the accuracy of the gLV suggests that in this setting continuous tuning of the environment yields a smooth slide of environmentally-relevant growth rates and negative pressures that directly produce a smooth variation of the stable fixed point in the full community.

### Environment-Induced Trends in Growth Rates and Interactions

Our fits and sensitivity analysis suggest that in the Glucose-variation setting a few interactions are responsible for the observed variation in dynamics and the shift in the equilibrium. Given the discussed nonlinearity of the loss function (Methods), stochastic optimization can yield solutions in a diversity of local minima with similar loss values. Thus, an ensemble of models - garnered from different runs of the optimization algorithm - is analyzed to identify key changes in the parameter set over the environmental gradients (Figure 4). The evolutionary algorithm employed tends to find similar local minima under fixed regularization parameters but is sensitive to the wander penalty. The wander penalty acts as an L1 sparsity regularization on the change in parameter set and, thus, the higher the weight, the more sparse the vector defined by the difference in any two sets. Under the largest penalties, parsimony in the variation highlights the importance of the interactions PK to RA (*α*_14_), RA to PA (*α*_61_), and PA to PK (*α*_46_) to explain the overall change in dynamics. These results appropriately match the set of parameters with the highest Sobol sensitivities (Figure S14, S15). We note that the lowest wander weight penalty yielded the fits that best generalized to the full community (lowest 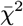) implying that it is more likely that a non-sparse ecological variance occurred and while a few key interactions varied significantly, the interaction between several organisms changed by a small amount as well.

**Figure 4:**
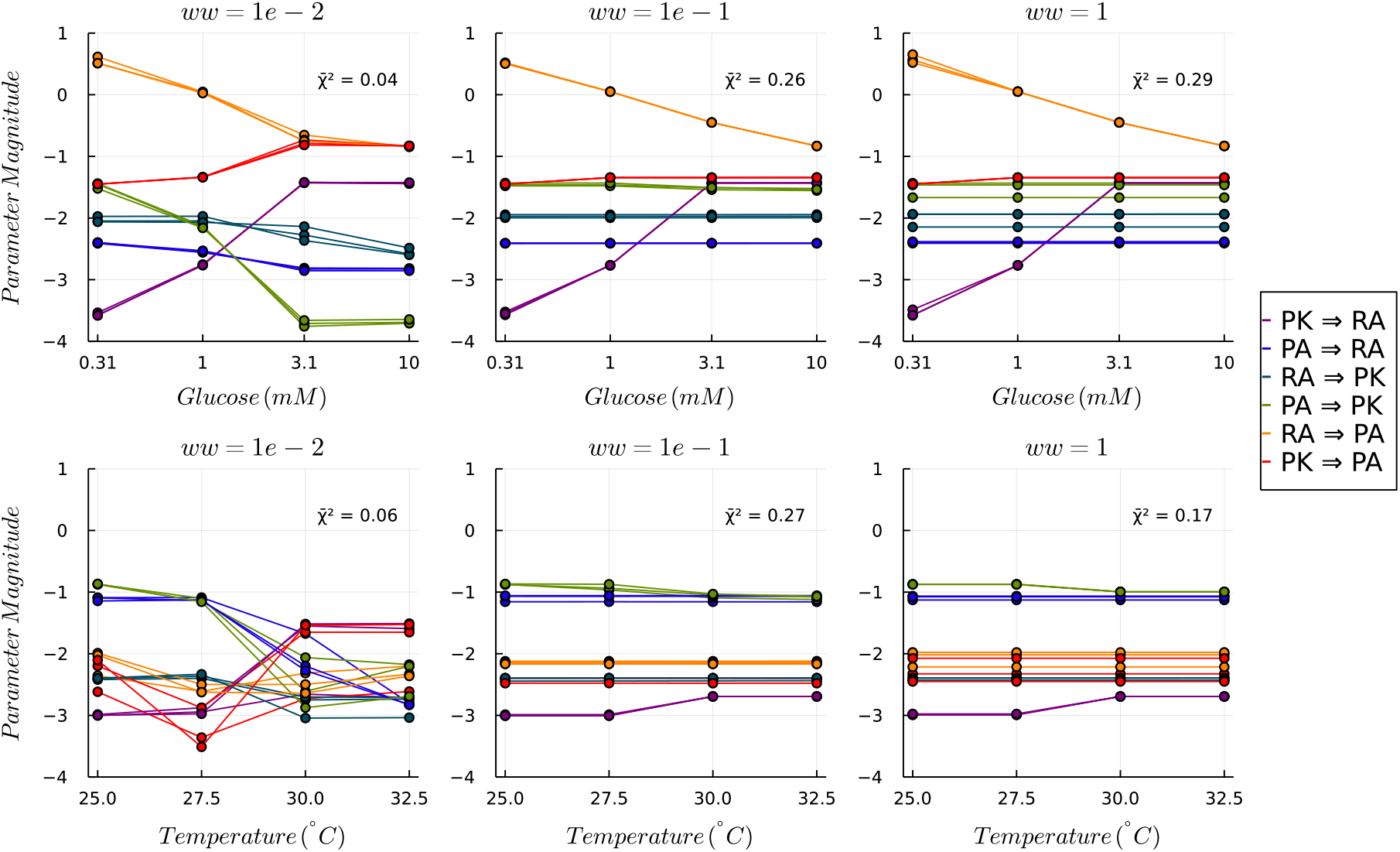
Principle Interaction Variance over Model Ensemble. Three parameter outputs of principle inter-species relationships are displayed for three different wander regularization penalties for both glucose and temperature settings. Mean fit score of ensemble, 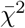 listed for each ww.

Interactions became more antagonistic as glucose concentration or temperature rose contrary to our initial hypothesis. In the principal subcommunity of RA, PK and PA, mean interaction decreases (but mean growth rate increases) as both glucose and temperature gradients increased (Table S3). While traditional ecological theory suggests that organisms are most amicable in “rich” resource settings [21, 22], in this community, the best-fit model suggests competitive interactions between members became stronger as one member, PA, began to take larger footholds in the community. We note that in the isogenic setting PA grows to the highest abundance of any species at the higher glucose concentration but only achieves a fraction of that in the corresponding setting when RA and PK are present. Hence, while glucose was more readily available for all organisms, this favored PA allowing it to grow significantly better, which actually reduced ecological gratuity in our fixed-capacity environment we hypothesize, thus, inducing hostility between species. We note that in previous works, others have demonstrated that competitive interactions stabilize fixed points with higher diversity [27], however, we observed that while this is true at low glucose and temperature regimes, eventually competitive interactions were dominated by the change in independent growth rates and self-interactions, allowing isogenic or nearly isogenic fixed points to dominate. This underscores two important ecological principles: firstly, that changes in inter-species interactions often correspond with changes in a population’s individual relationship with its environment [21], and that ‘starving’ a community can sum to positive ecological benefits that yield more desirable community dynamics [20, 21], as controller results will echo.

### Different Environments Yield Community Dynamics with Different Stability

The work here demonstrates that a change in the environment greatly influences the location and stability of the full community equilibrium, as observed other settings [20]. The microbiome at hand rapidly converges to the three dimensional subspace of *PA, PK* and *RA*, however, the 120-hour final point of this subsystem shifts from a rich diversity at low glucose and temperature to a *PA*-dominated system at high glucose and temperature [Figure 3]. The best-fit model suggests that the systems are indeed converging to fixed points with varying diversity as well as varying stability, which converge to cyclic hybrid trajectories that approach these fixed points before being reset upon passage (Figure S16). Interestingly, the best-fit model predicts that several of these cyclic hybrid trajectories capture unstable subcommunities in loops and otherwise converge to isogenic systems if simulated without dilution (Figure S16). The motion of trajectories in these scenarios is due to the dominating fixed point when left without passaging. No bifurcation in stability occurs over the change in environments, however, for initial conditions of low, equal abundance, the dominating fixed point moves from the species subspace of four or five members (with two of low abundance) to either two or single species subspaces (16).

The maximum real component of the eigenvalues of the Jacobian at each of these equilibria is studied to investigate the rate of convergence of the system - a common measure of stability [28] - near the fixed point at each environmental condition (Figure 5). Over the ensemble of fits corresponding to the lowest mean loss, the maximum eigenvalue falls indicating that the rate of convergence is faster for both higher glucose and temperature. Hence, for community trajectories near the fixed point, lower glucose and temperature systems will take more passages (or potentially seasons) to converge to a stable, cyclic hybrid trajectory. This is demonstrated in the virtual extended-passaging experiments (16). Variation in the rate of convergence implies that by manipulating the environment, it is possible to quicken or delay the system to potentially find dynamics more amenable to control. Generally, systems which evolve more quickly due to larger acceleration magnitudes are more difficult to manipulate, however, there might also be situations where we would like to accelerate the trajectory. This finding demonstrates that at the least one should consider the environment of their microbiome and how it might be perturbed, and generally that there are useful reasons to incorporate environmental control into microbiome therapies. As demonstrated later, the variation in rates of convergence is the foundation of the success of environment-varying control.

**Figure 5:**
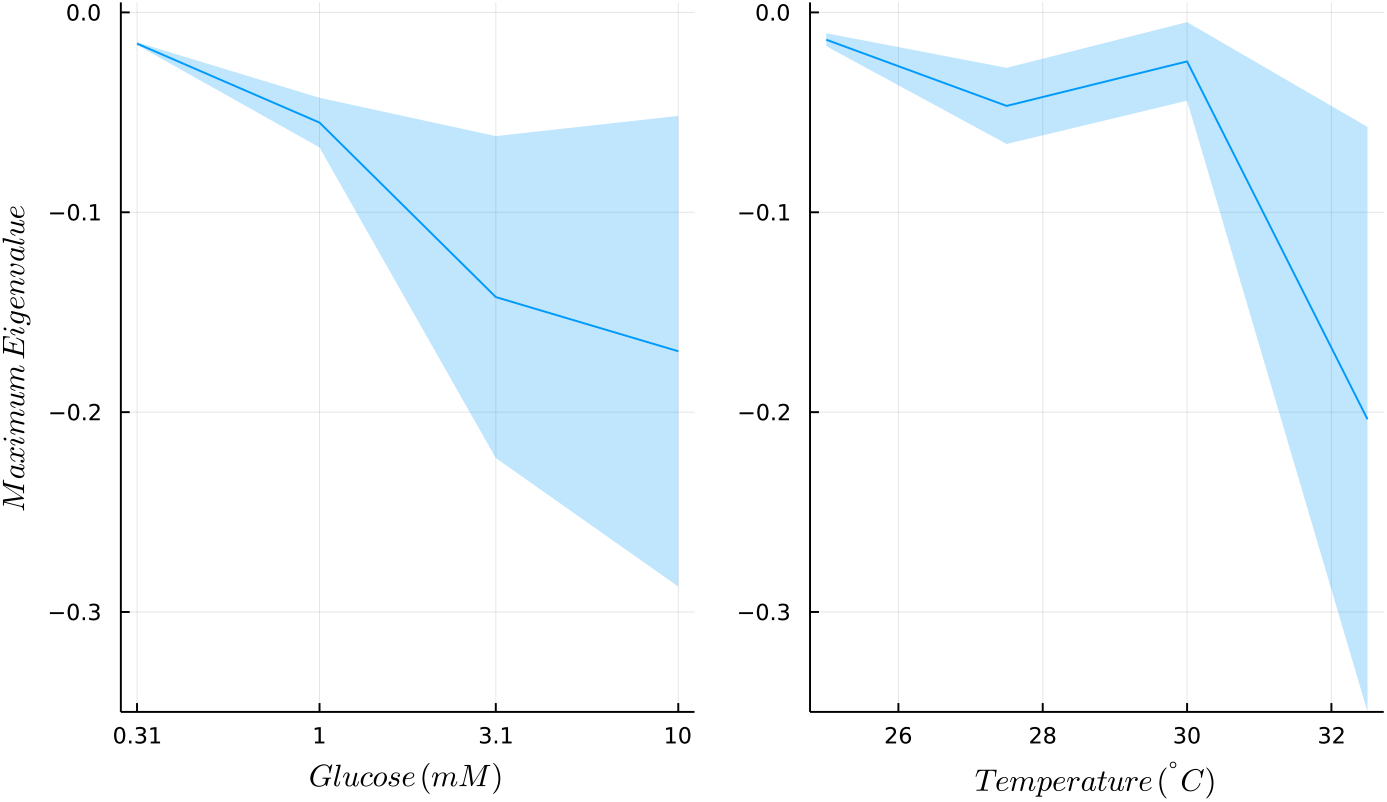
Rate of Convergence over Environmental Gradients. Mean maximum eigenvalue over the model ensemble is plotted for each glucose and temperature condition with band width determined by standard deviation.

### Theoretical Control of Environment-varying gLV Systems

With the parameterized microbiome model, we consider the problem of controlling a gLV system *in silico* with and without environmental variation. *In silico*, we consider the dynamics of only the 3 most abundant members with consistent representation in the diversity *in vitro* (with parameters in Table S3). The potential to control both temperature and glucose manifests through autonomous variation of growth rates *r* and interactions *A* and is analyzed in the context of driving the microbiome with minimal input to the following subcommunity equilibria: the *PA-PK-RA* fixed point, *PA-PK* fixed point, *PARA* fixed point, and the *PK-RA* fixed point. While the data show that one of these equilibria are stable, we include these targets to gauge how environmental control can slow or quicken convergence. For all problem formulations, we bound the input and disturbance to 16% and 8% of the observed maximum abundance to mimic a recalcitrant system or cautious program, and consider all possible input configurations (control of one, two, or all three members).

The overall objective of the optimal control programs is to drive the community as close as possible with minimal input, and we additionally quantify the number of states for which the target is achievable, defined as the controllable set [26]. This problem is inspired by the seasonal or situational need to select for different microbial compositions attributed to certain medical or agricultural benefits to the host.

We both simulate the forward perspective of driving the microbiome with a linearized Model-Predictive controller [25] from one equilibrium to another, and we compute and analyze the Backwards Reachable Tube (BRT) [26] for a given equilibria in time *τ* with a different Hamilton-Jacobi-Isaacs controller. The former is used for analyzing a continuous temperature model and the latter for a hybrid glucose model. The linearized MPC is a fast algorithm which has proved successful in the true nonlinear gLV dynamics [18], but stochasticity can impede success and MPC methods for nonlinear, hybrid systems are limited. Backwards-reachability is valuable for determining the boundary set of states which can be driven to an equilibrium under true stochastic, nonlinear dynamics, however, as a dynamic programming method it suffers in high-dimensional systems. Therefore, in the BRT analysis, we are restricted to considering limited schedules of hybrid-state switching and only the case of driving to the 3-membered equilibrium. However, this single situation allows us to explore how minimal and discrete environmental control, a more realistic option, can expand the set of controllable states. We demonstrate each as alternative approaches to controlling a microbiome when faced with different constraints.

Despite that changes in population dynamics appear similar for both environmental gradients, we assume that the underlying signals themselves differ such that two models for combining environmental data with distinct structures are necessary, one continuous and one hybrid. With respect to the problem of fitting nonlinear gLV systems, one might also choose a hybrid formulation because it restricts control to parameter domains where the model was trained unlike the continuous formulation which requires interpolation between fits. Ultimately, the two methods are demonstrated to offer different control approaches to various types of environment-varying factors.

### MPC with Continuous Temperature Control can Reduce Necessary Species Input

From an engineer’s perspective, temperature is an easily controlled variable. It is relatively unperturbed by the microbiome state itself and can often be manipulated externally with accuracy. Importantly, it remained constant in the parameterization experiments. Imagine that for these reasons, polynomial regression can capture the chemical changes that cause parameter changes in the experimented window of temperature change. This allows us to consider the problem of driving microbiomes in temperature-varying models with frequent manipulation of temperature (continuously or with Δ*t* = 0.1 intervals). We assume that the dynamics of stabilizing heat loss to a specific temperature are rapid and negligible compared to the gLV timescale (hours), thus the temperature-extended gLV system takes the form,

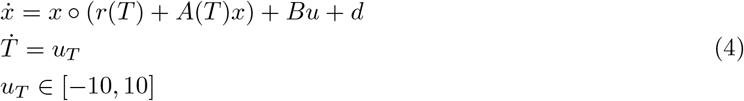

where *r*(*T*) and *A*(*T*) are the polynomial regressions of the temperature-parameter mapping and *u*_*T*_ is temperature input. *u*_*T*_ is bounded such that we might raise the temperature 2.5^*°*^*C* in 0.25 hours.

The general quadratic MPC program is used [25] with the linearized dynamics of the gLV-Temperature extended system with augmented state and input vectors 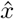 and *û*. Performance of the MPC is scored on two metrics: final species distance (the L2 distance between the final and target states after 6 hours of MPC action) and normalized total species input (the absolute sum of probiotic/antibiotic input during the program normalized by the number of species in the input configuration).

To explore this control scheme and the potential to modify temperature, the MPC is made to drive the state from each of the subcommunity equilibria to the others ten times, perturbed by random disturbances on each input (Figure 6). For each equilibrium-to-equilibrium path, all possible control species sets are tested and the depicted final species distance and total species input are averaged over all runs and sets.

**Figure 6:**
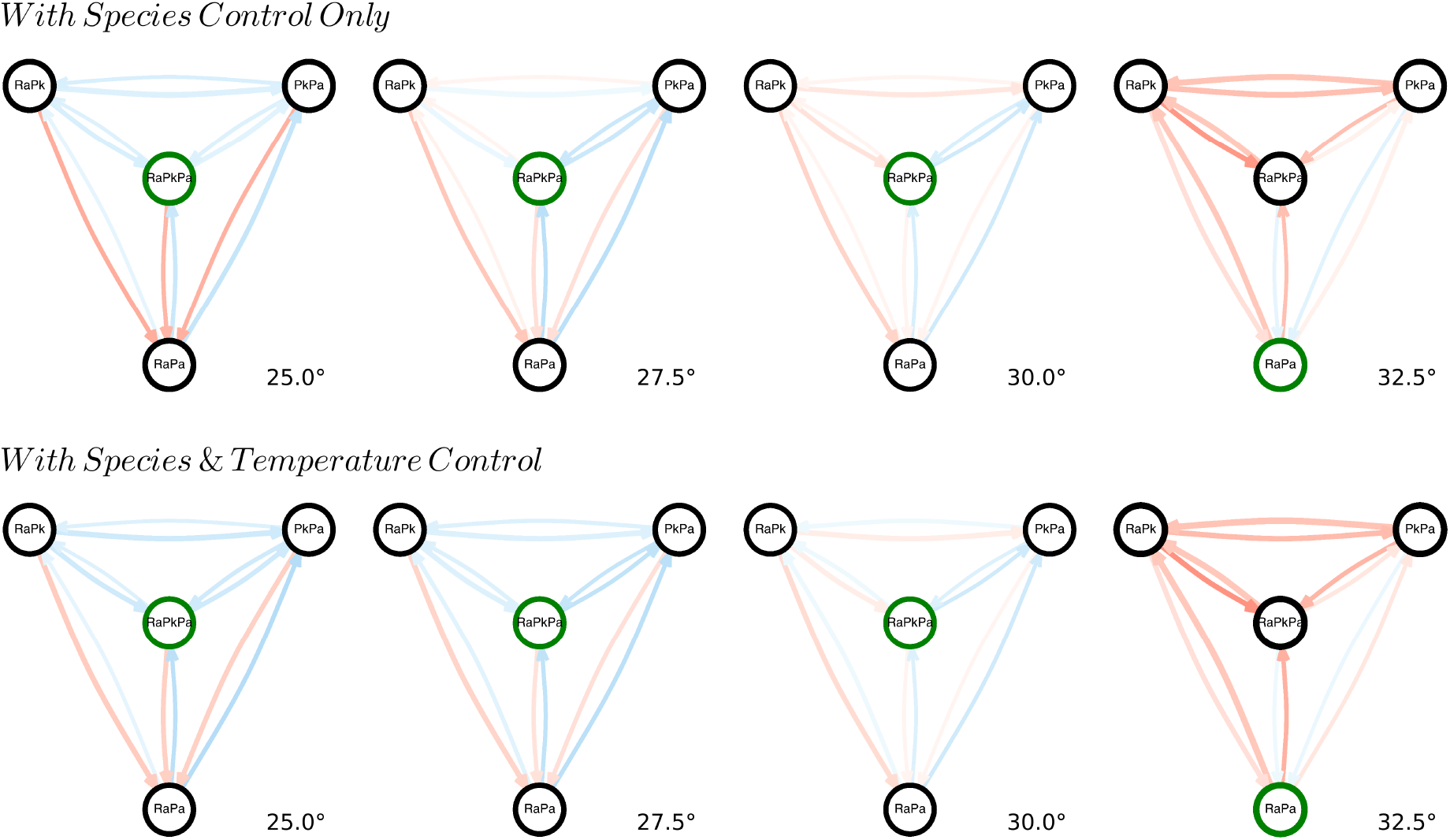
Path Success of MPC. Directed graphs depicting the initial-to-target paths on which the MPC was tested. The final distance to target determines the edge color (blue to red, ascending) and total species input determines the edge width of the linearized MPC. The data for each path is averaged over all possible control species sets. The stable equilibrium corresponds to the green node.

In both temperature controlled and fixed settings, the MPC is able to achieve closer final points with less input at lower temperatures (than higher) on most paths regardless of input configuration. There are some exceptions such as when the MPC is driven to the RA-PA equilibria which the model depicts as strongly unstable at lower temperatures but becomes the dominating autonomous fixed point at 32.5 degrees celsius (as seen in the experiments).

When averaging over all input configurations, across path-to-path variation the effect of temperature control offers minor differences for the MPC suggesting that minimal changes in limited configurations dominate the mean score. While there are significant improvements in paths to one state (RAPK) in intermediate temperatures, in general, the ability to transiently change the temperature in planning does not recover states that are poorly driven by the nominal controller on the given time span and planning horizon *for any input configuration*. We hypothesize this is due to the complexity of problems with limited configurations (e.g. control of only minor species RA) and the short comings of linearization in these harder problems, but we leave this for future work solely dedicated to mastering control of the microbiome.

For an input-driven understanding of the effect of temperature control, the same performance of the MPC with a given control species set is averaged over all paths (Figure 7). Grouping by input configuration reveals that the more species that can be actuated, the better the system is controlled with closer final distances and more efficient input usage. Regardless of configuration, the linearized system always proves controllable in the formal definition and in the nonlinear case, the network always needs only one minimal driver species to be structurally controllable (Not shown). Clearly, this belies variations controller performance, thus, while it might always be possible to drive to any point in the state space (definition of controllable), there is significant difference in the set of inputs when maneuvering the nonlinear space.

**Figure 7:**
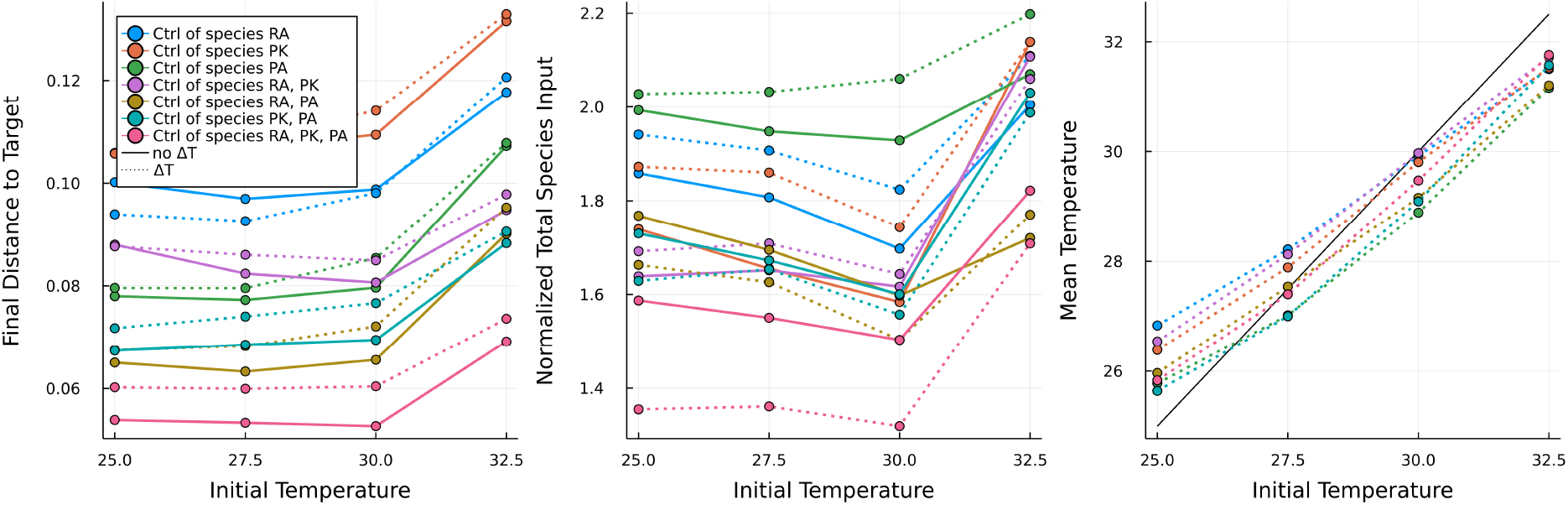
Summarized Success of MPC. Path and replicated averaged comparison of the linearized MPC with (dashed line) and without (solid line) the control of temperature for various sets of control species. (a) The final distance of the controller to the target equilibrium (b) The normalized, total species input in the control program (c) The mean temperature of the control program.

Notably, this analysis illuminates that if we are able to perturb PA and at least one other member, temperature control has an interesting ability to trade some final point accuracy for normalized total input; rapid temperature control allows pairwise manipulation of the system to become efficient without major detriments to accuracy. In general, the initial vs. mean temperature of the program reveals that the controller preferentially moves the system to intermediate temperatures to take advantage of faster dynamics that also support a rich diversity. Hence, the autonomous controller computes that optimal paths involve transiently perturbing the environment and performance indicates this is true.

With respect to future improvements, it is important to note that the incorporation of temperature control does not force the MPC to change the temperature and the trivial option of no temperature change is within the feasible set. Therefore, the deviation of the MPC with temperature control over the given horizon must have occurred because a change in temperature lowered the balanced cost between distance to target and sum input (Equation 11). This suggests the trade off between final species distance and sum input in these experiments is a result of the relative sizes of the penalty matrices *Q*, and *R* which could be altered to suit the designer.

In cases, such as when only one species was actuated, where adding temperature control resulted in both a higher final species distance and increased sum input, the temperature controller truly failed. Given that the trivial option of static temperature was considered, this failure can be traced back to two possible sources. First, there is the discrepancy between the linearized dynamics and the true nonlinear, stochastic gLV system, and its possible that over the horizon this error was exacerbated by the linearization of the polynomial, temperature-parameter relationship. Second, the horizon size of the MPC (*n* = 5) could’ve steered towards temperature domains yielding only short-term payoff, particularly in the cautious problem outlined above with small input constraints. While raising the horizon comes at a computational cost, the slow dynamics of the gLV system would likely allow a user to exert grander search and optimization efforts in online action, ameliorating both issues in future work.

Overall, the MPC simulations demonstrated the usage of temperature introduced freedom in the modulation of the microbial community to reduce the necessary quantity of probiotic and antibiotic input by approximately 20% at best at a sacrifice of 8% final distance to target at worst. It is important to note this was only available when actuating more than one species. Nonetheless, this option is valuable considering that probiotic and antibiotic inputs are expensive and can burden a host. This benefit may become more valuable if longer durations at these equilibria were required because the sum input will increase despite final distance remaining constant for the equilibria that are autonomously unstable (the majority). With high-throughput refinement of these control methods, we expect improved outcomes of observed results underscoring the value of environmental control in therapeutic control programs.

### HJI with Hybrid Glucose Control can Increase the Set of Controllable Community States

Glucose is a complex environmental signal. In batch systems, as populations grow, the concentration of glucose falls in proportion to each population. In such a system, the glucose dependent parameters are functions of the total abundance and diversity. The fit proves that constant gLV parameters reasonably predict the dynamics on multi-hour intervals (Figure 3) ie. that ecological pressures remain constant in this temporal window, perhaps because of sensing delays facilitated by kinetic thresholds. However, to assume this would remain true for rapid variations could be erroneous. If the interactions were majorly determined by delayed enzymatic repression and activation, sudden rises and falls in glucose concentration could induce more complex behavior, better represented by Hill terms rather than polynomials. Imagine that for these reasons, the problem of driving microbiomes in glucose-varying models takes a hybrid formulation [24, 29] (Figure 8), where each discrete “*q*-state” corresponds to a different set of parameters determined by the level of glucose introduced at restricted intervals. In addition to continuous state inputs *u*, we allow the discrete action *σ* of switching glucose-associated modes on 4-hour intervals. Each state shares the domain of the original model but unlike the experiments, the reset *R*_*σ*_ mapping associated with discrete state switching is the identity map (population dilutions might be unavailable in hydroponic culturing).

**Figure 8:**
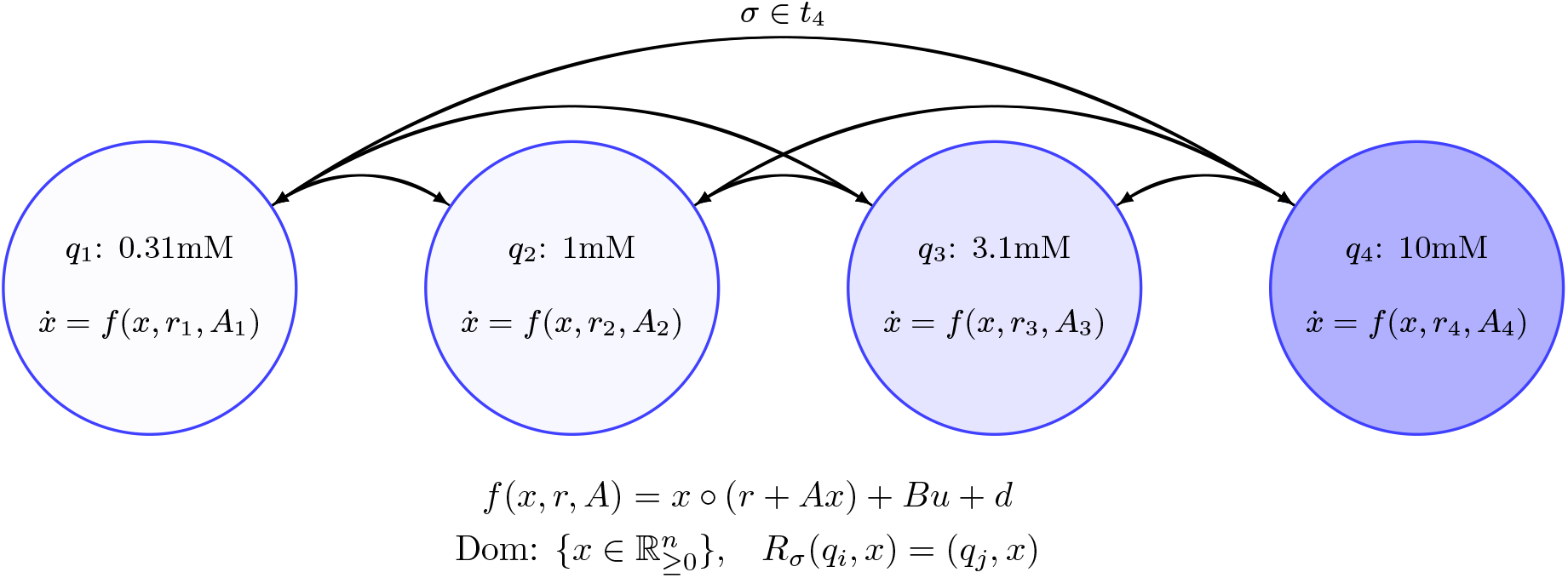
Definition of the Hybrid gLV System where discrete q-states correspond to different glucose concentrations and the corresponding gLV parameters for those concentrations. The domain *Dom* for all q-states is the zero-bounded state space. This system can be actuated by direct species input *u* or by changing the q-state via reset *R*_*σ*_ on 4-hour intervals.

We now define the optimal control problem as a zero-sum game between the controller and disturbance subject to the system dynamics, as is standard in HJI controller formulation [26]. The value function *J*(*x, t*) of the game is determined by proximity to the target equilibrium and thus, the controller is minimizing the distance to the equilibrium 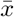 and takes the action for the worst possible disturbance. Given a window *t* ∈ [*τ*, 0], we compute the initial set of states *𝒢*(*τ*) for which the controller can safely drive to within an *ϵ* radius of 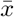 by *t* = 0 also known as the Backwards Reachable Tube (BRT)[26] or the set of controllable states (under the specific problem formulation). We use the volume of the BRT as a performance metric to compare the HJI controller with and without the ability to switch the environment.

We compute and compare the BRT of the RA-PK-PA equilibrium under fixed environments with environments subjected to scheduled switching. While the data showed this equilibrium is autonomously stable, we include this target to gauge how environmental control can change convergence and the set of controllable states. For the static environment (where no glucose state switching is allowed), we compute the BRT with *τ* = *−*12 hours for each of the four hybrid states corresponding to the initial glucose concentrations, and in the switched environment, we iteratively compute the BRT of *τ* = *−*4 hour segments in which the target set becomes the previous BRT computation, for all 4^3^ possible switch schedules.

The comparison of controllable state volumes for static and hybrid glucose environments is plotted for all control configurations (Figure 9). For some input configurations, including all two member control species sets, the computed BRT is small and minimally variant with switched or static environments suggesting uncompromising dynamics. Alternatively, in the case with control over only species RA, the entire searched state space falls within the BRT, and we are able to achieve sufficiently low distance to the goal state with or without switched environments. Finally, for some input configurations such as those with PK or all three species, the volume of controllable states under the fixed duration is increased. Generally, this reiterates the trend in the MPC simulations that the fully actuated system has the largest performance increase with environmental modulation. While expensive in practice, this further incentives the need for molecular or genetic tools to independently manipulate species in a microbiome. We can interrogate this performance-increase phenomenon through direct observation of the BRT surfaces: for each glucose concentration and it’s corresponding RA-PK-PA equilibrium, the static environment BRT and the maximum volume BRT of all possible switched schedules are juxtaposed when all species are controllable (Figure 10).

**Figure 9:**
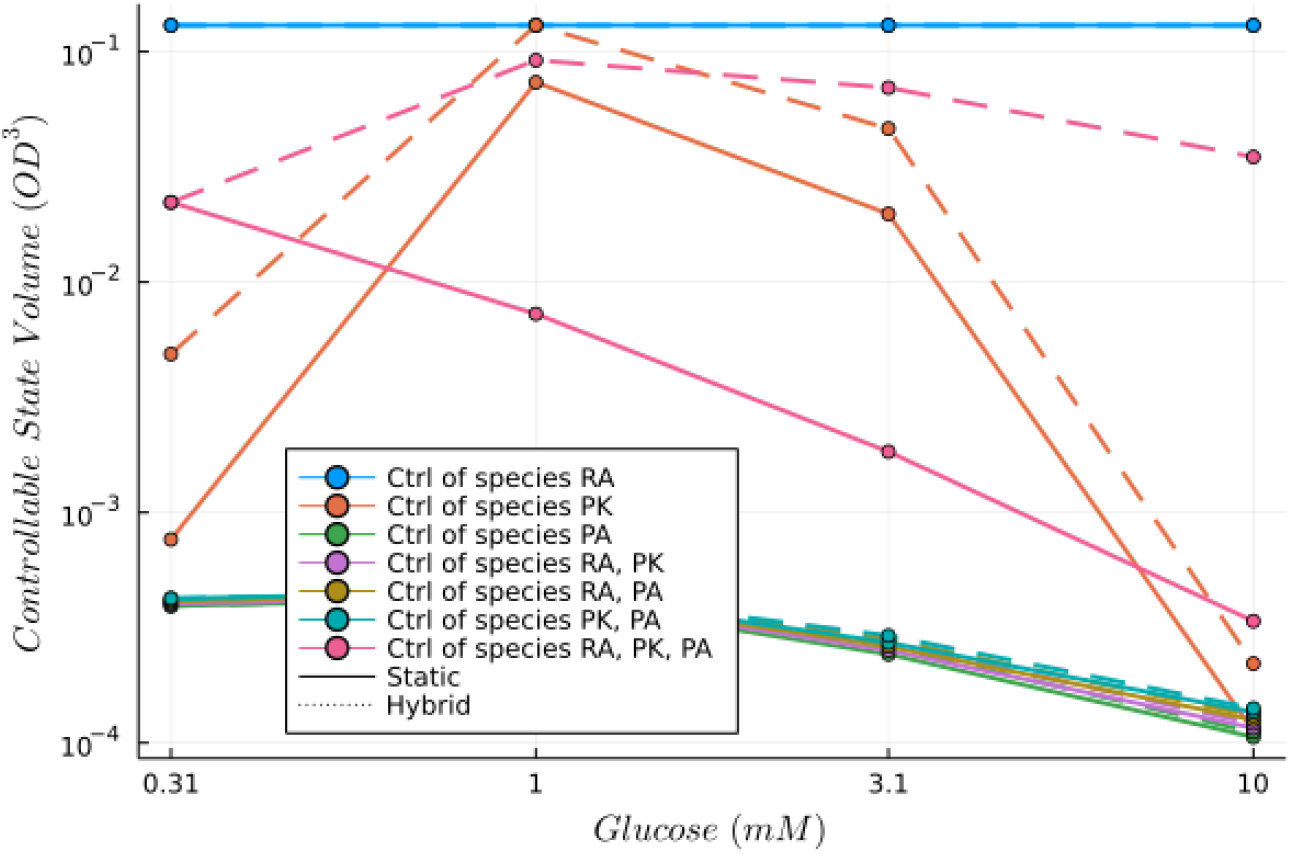
Comparison of BRT Volumes for Static and Switched Environments. The computed controllable state volume of the static environment problems (solid line) along with the hybrid schedule resulting in maximum volume (dashed line) for each of the glucose concentrations.

**Figure 10:**
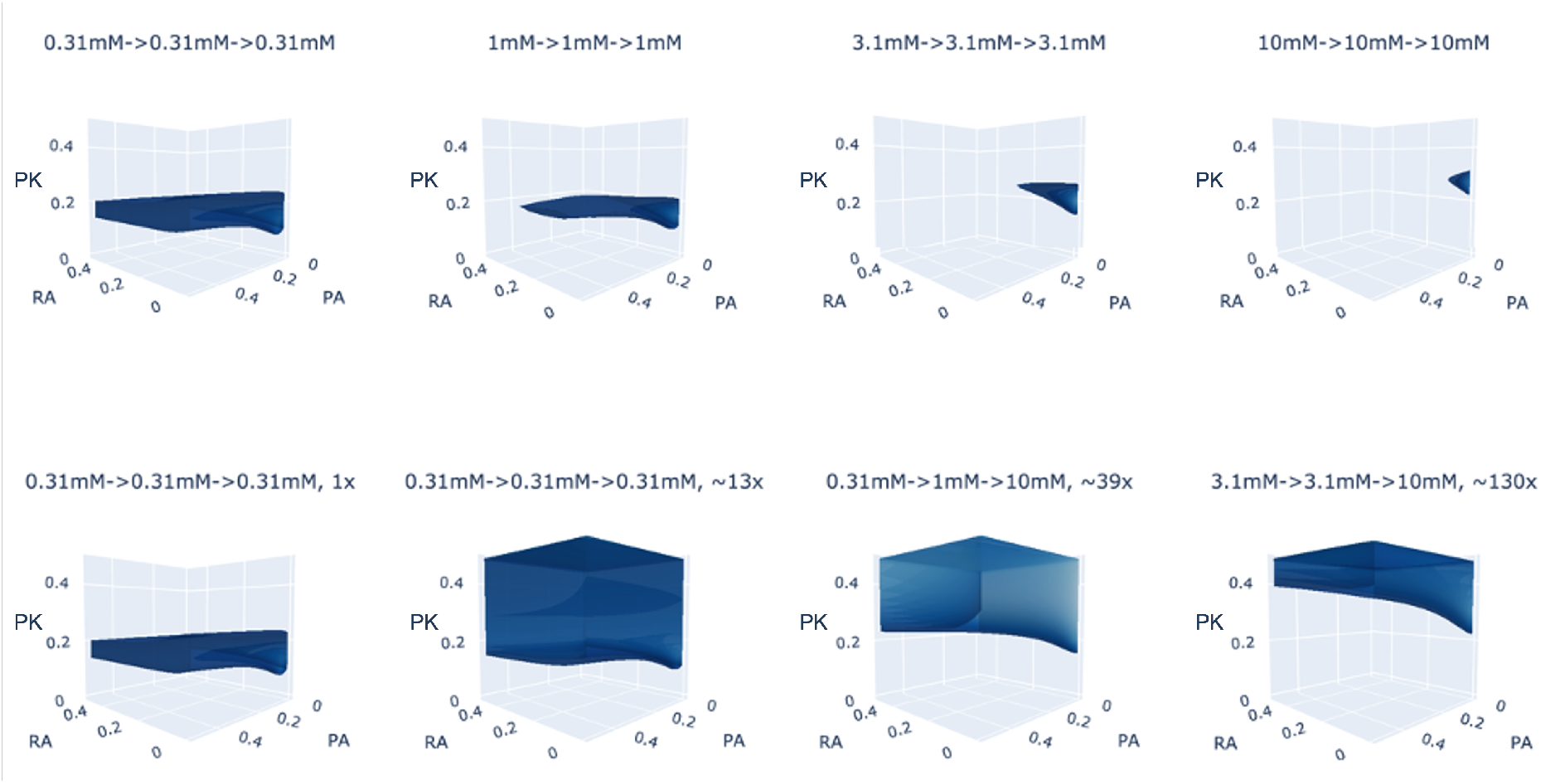
Backwards Reachable Tubes of Hyrbid Glucose HJI Control. Each column corresponds to the comparison of the fixed environment BRT (top) and maximum hybrid BRT (bottom) for driving to the fixed environment’s RA-PK-PA equilibrium. In this computation, the HJI controller can actuate all three species. The relative size of the maximum volume is listed in its title.

**Figure 11:**
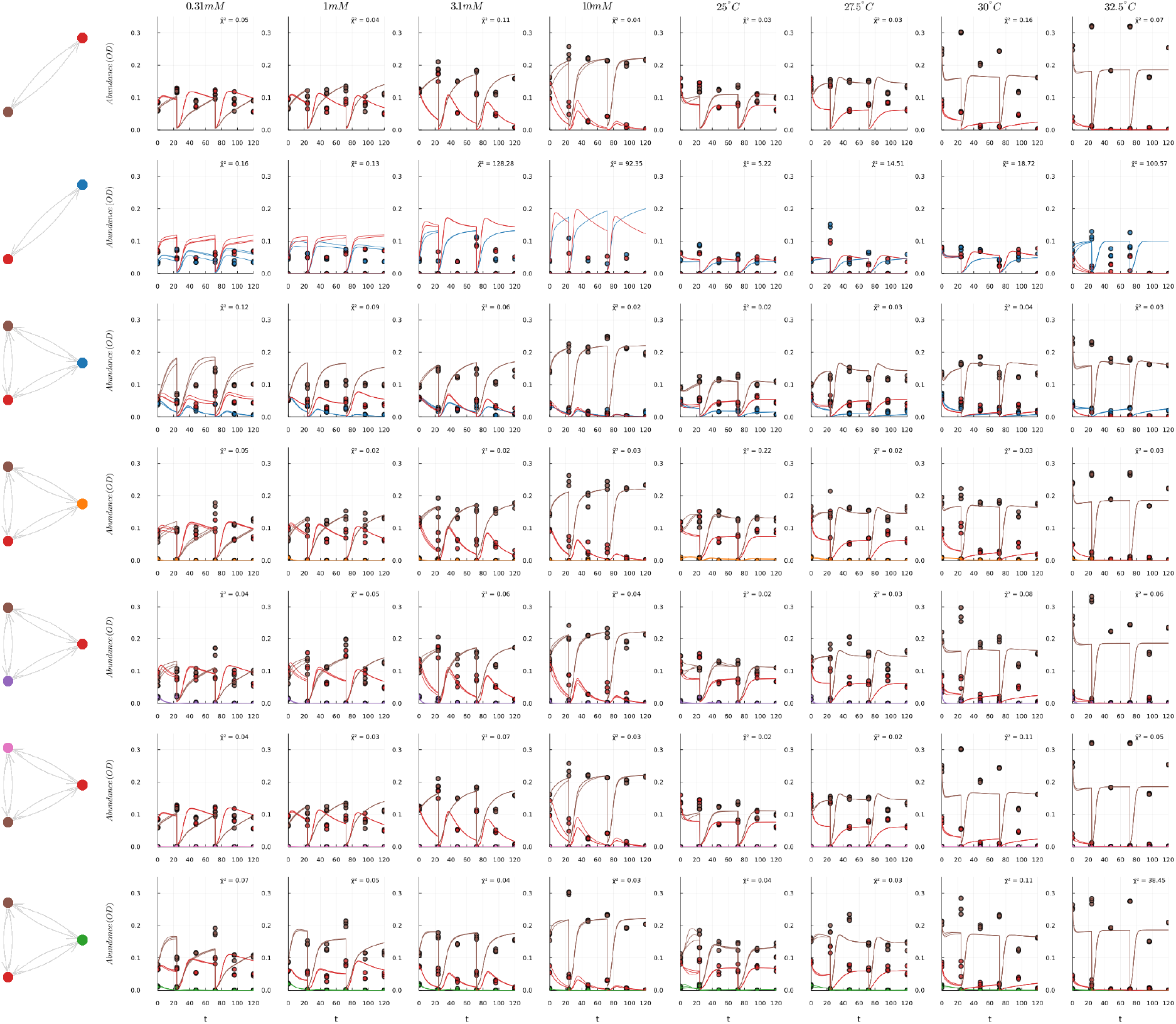
Model Fit on Subcommunity Training Data. The best fit compared with the training data of all subcommunity cultures. Real timecourse data are depicted as circles and model prediction is depicted as continuous lines.

As the static environment is the trivial schedule of switching to itself, the maximum BRT must be greater than or equal to the static BRT, however, we remarkably observe one instance of a BRT volume increase of 130*x* over a static environment when incorporating optimal glucose environment switching (Figure 10). In all cases, the maximum volume switched BRT includes utilizing the dynamics of lower glucose systems for at least two of the switches despite the rate of convergence analyses which revealed these systems to have the slowest rates of convergence. We interpret this outcome as a product of more efficient and effective input due to more amenable dynamics in the lower initial glucose concentrations which yield states with more diversity. When the glucose concentration is lower, attenuated growth of PA in particular allows for the presence of RA and PK which can then be used to influence one another by the control program, allowing the freedom to traverse a greater range of community states.

### Comparing MPC and HJI Behavior Illuminates the Utility of Different Environments

We note that the result of the hybrid HJI controller, to preference lower glucose systems, is slightly different than when continuous interventions were possible with the MPC, which suggested that intermediate temperatures allowed for highest accuracy and least input. If we ignore the difference in the two gLV models’ dynamics (which are relatively similar), we note that the controllers have slightly different priorities and operate at different intervals. The decision function of the HJI controller is irrespective of input magnitude, unlike the MPC, and seeks to drive the system to the target region against all random disturbance. In the lower glucose concentration environments, the higher average relative abundance allows for more input potential with farther reaching effect given that larger populations have greater influence on other species. Additionally, by also having slower, less-responsive dynamics in the lower glucose systems, stochastic disturbance is more manageable, allowing for ‘safer’ environments for the HJI controller. In the MPC, when input quantity influences decision, we can see that it becomes desirable to temporarily shift to systems where dynamics were quicker. Furthermore, because the MPC could act on rapid intervals, it could do this briefly without exposing itself to long durations of less-’safe’ dynamics where as the HJI controller would’ve locked itself into a less-’safe’ glucose regime for a 4-hour hybrid switch. In conclusion, our analysis demonstrates that the optimal strategy against stochastic disturbance is to wield environments that induce richer diversities with slower dynamics when operating without minimal input as a driving constraint, but otherwise, brief exposures to environments with faster dynamics can reduce input without sacrificing target accuracy.

## Conclusion

The model accuracy (Figure 3) of the experimental work further supports the high-resolution parameterization method proposed by Venturelli et al. [15] and demonstrates that variations in these parameters due to the environment are also predictable with the gLV system. As was previously known, microbiomes are heavily dependent on their environment and this work demonstrated that their composition is malleable in predictable manners with well fit models and quantified deformations. Practically, this knowledge is valuable for assessing how a community, designed or natural, might fluctuate under perturbations of the environment due to season, stress or design, and which members might dominate the total abundance in such situations. Notably, our improved gLV modeling illuminated the variance in stability and rates of convergence in different environments which are important to consider when controlling a microbiome. In future work, we hope to interrogate how the observed parameter variations compare to those of other environmental signals and those of the same signals of glucose and temperature on other communities. We are unable to answer why glucose and temperature induced such similar variations of dynamics and would like to explore if there exists a general variational pattern under arbitrary stress for this and other microbial communities.

The theoretical control of parameter-varying gLV systems demonstrated that knowledge of environmental dependence can be valuable when modulating a microbiome. These simulations underscore that certain environments are more amenable to altering a community. Our experiments demonstrate that, if achieving a target state is the only goal, environments yielding highest diversity and slowest rates of convergence are optimal. However, if minimal input is required, brief exposures to environments with faster dynamics can be highly beneficial. In general, our analyses demonstrate that controlling the environment can improve modulation programs by finding more optimal paths to target microbiome states - in terms of final distance to target diversities and total input - and by increasing the set of controllable states under certain input configurations. More sophisticated algorithms are required to take advantage of variations in competition in limited input configurations, perhaps by advanced numerical methods or reinforcement learning [14], and we leave this to future work.

In summary, these experiments, in vitro and silico, demonstrated how environment-dependent competitive dynamics within the soil microbiome can be modeled by parameter-varying gLV systems and that this knowledge can be valuable for improving prediction and control of these systems. Both of these findings offer new directions for designing and controlling microbial communities in human health, agriculture and chemical production.

## Materials and Methods

### Isolation Protocol

Mature *o. sativa* were harvested preserving several inches of roots. The roots were placed in 25% glycerol, incubated for 20 minutes at room temperature and then on dry ice for 30 minutes. Afterwards, roots were vortexed and washed with sterile water twice and then sliced into 1cm sections. Next, root slices were deposited into 1X phosphate-buffered saline (1g roots/3mL PBS) that was poured into a sterile, ceramic mortar and ground modestly to release cells into the buffer solution. The buffer solution was then serially diluted from to 10^*−*1^ to 10^*−*7^ dilutions of which 1 *µ*L was used to inoculate 200 *µ*L of 10% TSB. After observing growth, DNA was extracted and amplified from the isolates via the procedure outlined below. Most-similar blast results and strain identification is listed in supplemental Table S1.

### Culture Methods

All organisms were grown to an OD of 0.2 before being combined in equal proportion to make the pairs, triplets, and full community cultures with a Biomek Liquid Handling Robot (model, part number, etc.). All cultures were grown in triplicate in 2mL of 10% TSB, inoculated to a combined OD of 0.001, and passaged after 48 hours with 1:20 dilution, twice (144 hours total). On 24 hour intervals, 200 uL was removed for OD reading and 400 uL was removed to spin down and storage at -20C prior to DNA extraction and sequencing (prior to passaging).

### DNA Extraction and Amplification

For cost and simplicity, the Direct PCR method [30] (Figure S12) was used to extract genomic DNA from the centrifuged biomass. Sample pellets were resuspended in the surfactant Igepal, 0.5% final concentration, then freeze-thaw cycled to lyse bacterial cells. The released DNA was used as template to amplify the V4/5 region of the 16s gene based on the Earth Microbiome Project [31-515F, 32-926R] and samples were tracked with a dual index strategy.

### Sequencing and Quantification

The amplicons were sequenced on an Illumina MiSeq with 600 bp v3 reagents and the sequencing results were processed with Perl scripts [33] to yield reads counts. Samples were demultiplexed using the index read of the forward read (IT001-IT096) with Illumina software then forward and reverse reads were merged with Pear [34]. Reads with more than one expected error were filtered out with Usearch [35] then demultiplexed with the reverse primer indices (inline1-inline96), and finally, Unoise3 [36] was used used to filter out rare reads and chimeras. A count table of each isolate per sample was made by matching sequences with original isolate 16S reads.

Relative abundances were computed from the counts and then multiplied with the corresponding OD sample to estimate each species’ absolute abundance. These computed absolute abundances comprised the time course data for each isogenic or subcommunity culture that was then used for training the gLV model.

### Experimental Design

Pairwise co-culture parameterization of community models results in high fidelity predictions of species growth [15, 16, 37, 38]. While the method can produce reliable models however, pairwise fitting is expensive. The number of pairs grows with *𝒪*(*n*^2^) and each pair requires data from multiple time-points, which need to be processed to identify diversity and total abundance. With prior knowledge of full-community diversity, it is possible to compromise experimental labor with model fidelity. In general, growth rates and interaction parameters in community dynamic models increase with the population size of their individual species. Thus, rare species often (but not always) have lesser influence on community dynamics, and interactions between them are less relevant. We used this principle to guide the selection of training data.

We fit the gLV model with data from seven isogenic cultures, two pairs between the three most abundant members, and from five triplicates between the two most abundant members and each of the other five members (Figure 1). This set allowed fitting of 64% of all parameters (seven growth rates *r*, seven self-interactions *α*_*ii*_, 22 interactions *α*_*ij*_) with 14 rather than 28 cultures (for 7 isogenic and all 21 pairwise cultures). The model is validated by measuring fit (defined by lowest *χ*^2^ loss [39]) to the growth of full community cultures in both static and switched environments where glucose concentration or temperature is changed on 48-hour schedules.

### Parameterization

The gLV model for each of the 8 conditions (4 glucose points, 4 temperature points) is trained via a two part ‘global-local’ optimization strategy. The fits that best predicted the full community data were trained with BlackBoxOptim’s adaptive differential evolutionary algorithm (ADE) [40], although several well-known gradient-based algorithms were tried. To counter the gLV’s hyper flexibility and elucidate the underlying ecological phenomenon, a penalty on the deviation between condition-adjacent fits was included to encourage a parsimonious difference in parameters (Eqn). With respect to the dynamics, the simulations are initialized from the first data point to negate the stochasticity of community establishment [41] and also include the discrete event of 1 : 20 passaging on 48 hour intervals.

The ‘global’ and ‘local’ optimization programs share the same loss function (Eqn 5) but are initialized from different positions in the state space. The ‘global’ leg is initialized with random values spanning fixed parameter ranges *µ* ∈ [0, 1], *α*_*ii*_ *∈* [*−*5, *−*0.5], and *α*_*ij*_ *∈* [*−*5, 5], but with initial growth rates and self interactions set to estimates made from endpoints of the single-species growth assays. The ‘local’ leg is initialized with the optimal solution of the global run and uses parameter ranges which are within 50% of the previously optimal vector. The loss function for both legs iterates over all subcommunity cultures (singles, pairs and triplets) to sum a total cost so that all parameters are optimized on the full training data set. This allows parameters which are utilized in multiple subcommunities to be fit to multiple experiments simultaneously. All parameterization scripts were written in Julia [42].

The loss is defined as the replicate-averaged, sum L2 norm between observed states *x*_*t*_ and gLV predicted 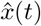 for time points *t ∈ t*_*p*_ with L1 regularization on the parameter set and between condition-adjacent parameter sets. Thus, the loss takes the form,

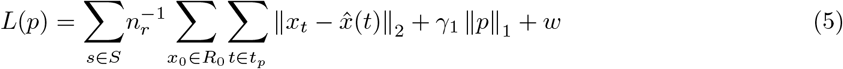

where *p* is the flattened vector of gLV parameters *r* and *A, S* is the set of all subcommunities, *R*_0_ is the set of initial states of each replicate, *n*_*r*_ is the number of replicates for each subcommunity (which varies because a few were contaminated and excluded), *γ*_1_ is the regularization coefficient, and *w* is the wander penalty. We chose to fit the lowest glucose and temperature condition with *w* = 0 and then for each increase in glucose or temperature *w* becomes the weighted norm between the parameters being fit *p* and the previous fit’s parameters *p*_*c*_,

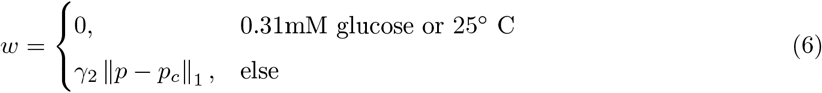

Hence, the loss encourages the fits of different environmental conditions to differ in a minimal number of parameters from one another.

The training data (points) and best model fit (smooth lines) are displayed in (11). In most subcommunity cultures, PK and PA dominate the absolute abundance (Figure 11), occupying *>* 90% of the relative abundance, despite significant growth in isogenic contexts by all species. We compute the Sobol Indices [43] of the optimal parameter sets with respect to the loss in order to verify the fit (Figures S14, S15).

Due to the flexibility of the gLV and the stochastic nature of evolutionary algorithms, a model ensemble is generated to strengthen our belief in trends observed across possible fits. The ensemble is generated for each of the environmental conditions from several runs of the same optimization hyperparameters. The “best-fit” model is the model of the ensemble that had the lowest final loss for all attempted hyperparameter values. To identify key variations in the parameters, the ensembles of several wander weight penalties are compared (Figure 4).

### Stability Analysis

The non-trivial equilibria of the autonomous gLV system satisfy either of the following equations,

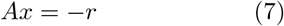

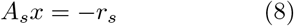

for any subcommunity *S* : [1, *n*] \*{i}* where *x*_*i*_ = 0. While full-community fixed points often include negative values for specific states, in the bounded system the trajectory is intercepted at the *x*_*i*_ = 0 subspaces resulting in a subcommunity without member(s) *i*. Furthermore, in systems with scheduled resets (eg. due to passaging), forced cycling [29, 44] may appear where trajectories are reset into regions of stability where they previously converged (Figure S16). As demonstrated by Gore et al., in the continuous case as well, dilution can similarly create alternative stable states [20]. Here we estimate the dominating equilibrium as that of the dominating subcommunity after simulating two passages and integrating the system trajectory for an additional 200 hours.

The maximum real component of the eigenvalues of the Jacobian at each of the dominating equilibria given by,

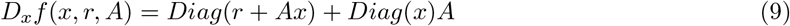

is analyzed in order to illuminate the rate of convergence of the system. This is done for each model in the ensemble corresponding to the lowest average loss and the mean and standard deviation of the maximum real components is compared to observe variation across the environmental gradients (Figure 5).

### Control Program Details

Species input and disturbance bounds are chosen to be 16% and 8% of the real maximum abundance (0.3),

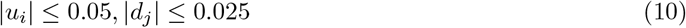

to simulate either a recalcitrant system or a cautious modulation program. For all control simulations, we neglect continuous variations of input sensitivities *B* but consider binary variations of it corresponding to different input configurations, eg. *B* equal to the identity matrix (control of all species) versus *B* equal to the matrix of all zeros except one in the uppermost left (control of RA only).

### MPC Methods

The MPC objective function penalizes the state and input paths, 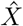 and *Û*, by the distance to the target point 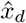 running state-cost matrix *Q* and running input-cost matrix *R*. Thus, for the full state vector 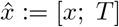 and input vector *û*:= [*u*; *u*_*T*_] with path matrices 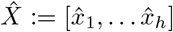 and *Û* := [*û*_1_, … *û*_*h*_], the nonlinear program seeks to minimize,

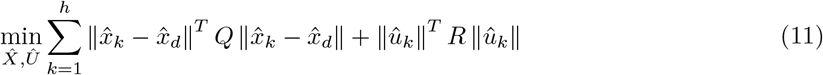

subject to the aforementioned state constraints and Forward Euler dynamics constraints,

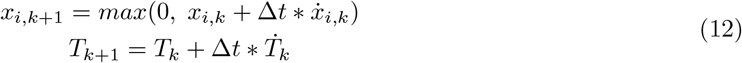

In practice, mapping the discrete algebraic Ricatti solution of the unconstrained problem to the above input bounds resulted in a controller that significantly outperformed the solutions produced by optimization algorithms with 1000x faster runtime.

For each of the MPC input-to-target paths, we randomly generated 10 vectors of disturbance which were added at each time step to test the controller. The same disturbance vectors were used for the MPC with and without the ability to modify temperature and performance metrics were averaged over all ten runs. The interpolations for growth rates and interactions in the continuous, Temperature-varying gLV model were computed with 4th-order polynomials in Julia which defaults to using the Gauss-Newton method.

### HJI Controller Methods

For the HJI controller, we define the optimal control problem as a zero-sum game between the controller and disturbance subject to the system dynamics, as is common in HJI controller formulation [26]. The value function *J*(*x, t*) of the game is determined by proximity to a target equilibrium 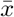,

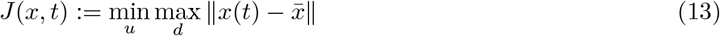

thus, the controller is minimizing the distance to the equilibrium and takes the action for the worst possible disturbance. Given a window *t* ∈ [*τ*, 0], we seek the initial set of states *𝒢*(*τ*) for which the controller can drive to within an *ϵ* radius of 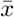 by *t* = 0,

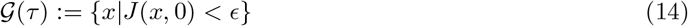

also known as the Backwards Reachable Tube (BRT)[26]. To compute the BRT, it is possible to solve the following Hamilton-Jacobi-Isaac’s PDE for which *J*(*x, t*) is a viscosity solution,

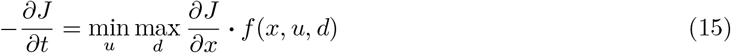

While the exact solution to this PDE is unknown in general, it is possible to obtain a numerical approximation for the level set of *G*(*τ*) by gridding a subspace of ℝ^*n*^ and recursively evaluating the change in *J* over time [26]. These computations are done with the Python toolbox OptimizedDP [45, 46].

This computation is done over a time span of *τ* = *−*12 for all possible environment switches after every 4 hours, including the trivial case of switching to the same environment. The volume of the surface at the end of the computation is approximated by summing the number of points in the grid satisfying the value function condition *J*(*x, τ*) *<*= 0 multiplied by the grid cell volume.

## Data, Code and Reagent/Strain Availability

All raw and processed data along with scripts used in the work have been published to the github repository TECM (https://github.com/willsharpless/TECM) with the exception of the Perl scripts used to process the illumina reads [33]. The raw reads were additionally published to the github under the “data/Raw/” directory. Isolate strains are available by request from the Arkin Lab Strain Collection.

## Author Contributions

WS conceived the project and refined the idea with APA and KS. WS, KS, FS, JK carried out the experimental work. WS wrote the manuscript and all authors contributed to editing.

## Acknowledgements

We would like to thank Dr. Devin Coleman-Derr and Dr. Alex Styer for supplying rice plants and ideas about cultivating the isolates. We would like to thank Dr. Adam Deutschbauer and Dr. Hans Carlson for their counsel and assistance in the pooled sequencing of all pairwise and triwise experiments. We would like to thank Morgan Price for his Perl scripts for analyzing the sequencing results. We would like to thank Dr. Claire Tomlin and Dr. Shankar Deka for valuable discussions on Model Predictive and Hamilton-Jacobi controllers.

## Notes

https://github.com/willsharpless/TECM

### Competing Interest Statement

The authors have declared no competing interest.

