## Supplemental for "Towards Environmental Control of Microbiomes"

| Isoalte abbrev. | Most sequence-similar organism | Sequence Identity | Rice Soil | Arkin Lab Strain ID |
|---|---|---|---|---|
| iSK | *Shinella kummerowiae* | 75.0% | Mojave Roots | APA7520 |
| MP | *Microbacterium phyllosphaerae* | 98.6% | Calcine Clay | APA7521 |
| PK | *Pseudomonas koreensis* | 98.1% | Regolith | APA7522 |
| BM | *Bacillus megaterium* | 99.5% | Mojave Roots | APA7523 |
| PA | *Pantoea agglomerans* | 98.6% | Calcince Clay | APA7524 |
| iFG | *Flavobacterium ginsengiterrae* | 83.5% | Calcine Clay | APA7525 |
| RA | *Rhizobium pusense/Agrobacterium salinitolerans* | 98.5% | Mojave Roots | APA7526 |

**Table 1: Strains**. Information about each strain used in the work including the most similar organism by sequence similarity, the corresponding sequence similarity, the soil in which the rice plant that yielded the organism was grown and the Arkin lab strain archive ID.

| | *r* | *A* | | | | | | |
|---|---|---|---|---|---|---|---|---|
| 0.31mM | 0.5 | -2.4 | 0.0 | 0.0 | -3.6 | 0.0 | -2.0 | 0.0 |
| | 0.2 | 0.0 | -0.6 | 0.0 | -2.7 | 0.0 | -2.2 | 0.0 |
| | 0.4 | 0.0 | 0.0 | -1.6 | -2.4 | 0.0 | -2.4 | 0.0 |
| | 0.4 | -2.1 | 2.5 | -2.0 | -2.8 | 3.4 | -1.5 | 1.8 |
| | 0.5 | 0.0 | 0.0 | 0.0 | -2.9 | -2.2 | -3.7 | 0.0 |
| | 0.3 | 0.5 | 4.5 | 4.4 | -1.4 | 4.9 | -1.1 | 1.2 |
| | 0.5 | 0.0 | 0.0 | 0.0 | -2.0 | 0.0 | -3.7 | -1.9 |
| 1mM | 0.5 | -2.3 | 0.0 | 0.0 | -2.8 | 0.0 | -2.5 | 0.0 |
| | 0.3 | 0.0 | -0.9 | 0.0 | -2.7 | 0.0 | -2.2 | 0.0 |
| | 0.4 | 0.0 | 0.0 | -1.7 | -2.4 | 0.0 | -2.4 | 0.0 |
| | 0.5 | -2.1 | 2.5 | -2.0 | -2.7 | 3.4 | -2.2 | 1.8 |
| | 0.5 | 0.0 | 0.0 | 0.0 | -2.9 | -1.9 | -3.7 | 0.0 |
| | 0.3 | 0.0 | 4.5 | 4.4 | -1.3 | 4.6 | -1.6 | 1.2 |
| | 0.5 | 0.0 | 0.0 | 0.0 | -2.0 | 0.0 | -3.6 | -1.9 |
| 3.1mM | 0.5 | -2.2 | 0.0 | 0.0 | -1.4 | 0.0 | -2.9 | 0.0 |
| | 0.3 | 0.0 | -1.1 | 0.0 | -2.7 | 0.0 | -2.2 | 0.0 |
| | 0.5 | 0.0 | 0.0 | -2.2 | -2.4 | 0.0 | -3.2 | 0.0 |
| | 0.6 | -2.1 | 2.5 | -2.0 | -2.4 | 3.4 | -3.7 | 1.9 |
| | 0.4 | 0.0 | 0.0 | 0.0 | -2.9 | -1.5 | -3.7 | 0.0 |
| | 0.3 | -0.8 | 4.5 | 4.4 | -0.8 | 4.6 | -1.8 | 1.2 |
| | 0.5 | 0.0 | 0.0 | 0.0 | -2.0 | 0.0 | -3.6 | -1.9 |
| 10mM | 0.6 | -2.1 | 0.0 | 0.0 | -1.4 | 0.0 | -2.9 | 0.0 |
| | 0.4 | 0.0 | -1.1 | 0.0 | -2.7 | 0.0 | -2.2 | 0.0 |
| | 0.6 | 0.0 | 0.0 | -2.4 | -2.4 | 0.0 | -3.2 | 0.0 |
| | 0.7 | -2.5 | 2.5 | -2.0 | -2.0 | 3.4 | -3.6 | 1.9 |
| | 0.4 | 0.0 | 0.0 | 0.0 | -2.9 | -1.5 | -3.7 | 0.0 |
| | 0.4 | -0.8 | 4.5 | 4.4 | -0.8 | 4.6 | -1.9 | 1.3 |
| | 0.4 | 0.0 | 0.0 | 0.0 | -2.0 | 0.0 | -3.6 | -1.6 |

| | *r* | *A* | | | | | | |
|---|---|---|---|---|---|---|---|---|
| 25C | 0.3 | -5.0 | 0.0 | 0.0 | -2.0 | 0.0 | -0.8 | 0.0 |
| | 0.4 | 0.0 | -1.1 | 0.0 | 0.3 | 0.0 | -2.9 | 0.0 |
| | 0.3 | 0.0 | 0.0 | -1.0 | 0.5 | 0.0 | -2.0 | 0.0 |
| | 0.3 | -4.8 | 1.7 | -4.9 | -3.3 | -4.9 | -0.6 | -0.4 |
| | 0.6 | 0.0 | 0.0 | 0.0 | -4.2 | -4.8 | -3.9 | 0.0 |
| | 0.3 | -3.6 | 4.9 | 4.9 | -2.3 | 5.0 | -1.3 | -0.2 |
| | 0.3 | 0.0 | 0.0 | 0.0 | 0.5 | 0.0 | -2.9 | -2.3 |
| 27.5C | 0.3 | -4.5 | 0.0 | 0.0 | -2.0 | 0.0 | -1.0 | 0.0 |
| | 0.4 | 0.0 | -1.3 | 0.0 | 0.2 | 0.0 | -2.7 | 0.0 |
| | 0.4 | 0.0 | 0.0 | -1.5 | 0.3 | 0.0 | -2.4 | 0.0 |
| | 0.4 | -4.5 | 1.9 | -4.0 | -3.9 | -4.1 | -1.0 | -0.4 |
| | 0.5 | 0.0 | 0.0 | 0.0 | -4.2 | -4.8 | -4.5 | 0.0 |
| | 0.6 | -3.4 | 4.0 | 4.5 | -2.5 | 4.8 | -3.1 | -1.2 |
| | 0.3 | 0.0 | 0.0 | 0.0 | 0.3 | 0.0 | -3.5 | -2.6 |
| 30C | 0.3 | -4.4 | 0.0 | 0.0 | -1.8 | 0.0 | -1.5 | 0.0 |
| | 0.5 | 0.0 | -1.4 | 0.0 | 0.6 | 0.0 | -2.7 | 0.0 |
| | 0.4 | 0.0 | 0.0 | -1.8 | 0.6 | 0.0 | -2.6 | 0.0 |
| | 0.4 | -4.4 | 1.8 | -3.9 | -3.7 | -4.3 | -2.0 | -0.6 |
| | 0.3 | 0.0 | 0.0 | 0.0 | -4.5 | -4.7 | -4.0 | 0.0 |
| | 0.6 | -3.3 | 4.2 | 4.9 | -1.8 | 4.9 | -3.5 | -1.0 |
| | 0.6 | 0.0 | 0.0 | 0.0 | 0.2 | 0.0 | -3.7 | -2.5 |
| 32.5C | 0.4 | -4.2 | 0.0 | 0.0 | -1.9 | 0.0 | -1.9 | 0.0 |
| | 0.3 | 0.0 | -1.3 | 0.0 | -0.4 | 0.0 | -2.7 | 0.0 |
| | 0.3 | 0.0 | 0.0 | -1.9 | 1.0 | 0.0 | -2.4 | 0.0 |
| | 0.3 | -4.1 | 1.2 | -3.9 | -3.6 | -4.3 | -1.8 | -0.5 |
| | 0.4 | 0.0 | 0.0 | 0.0 | -4.6 | -4.9 | -3.7 | 0.0 |
| | 0.6 | -3.3 | 4.0 | 4.7 | -1.8 | 4.8 | -3.4 | -1.3 |
| | 0.6 | 0.0 | 0.0 | 0.0 | 0.2 | 0.0 | -4.0 | -2.7 |

**Table 2: Parameters of the Best Fit Model** with optimization hyper-parameters $\lambda_1 = 1e-4$, $\lambda_2 = ww = 1e-2$, $r \in [0.0, 1.0]$, $\alpha_{ij} \in [-5.0, 5.0]$. Data stored in *p_full_all.jld* file on github repository TECM (https://github.com/willsharpless/TECM)

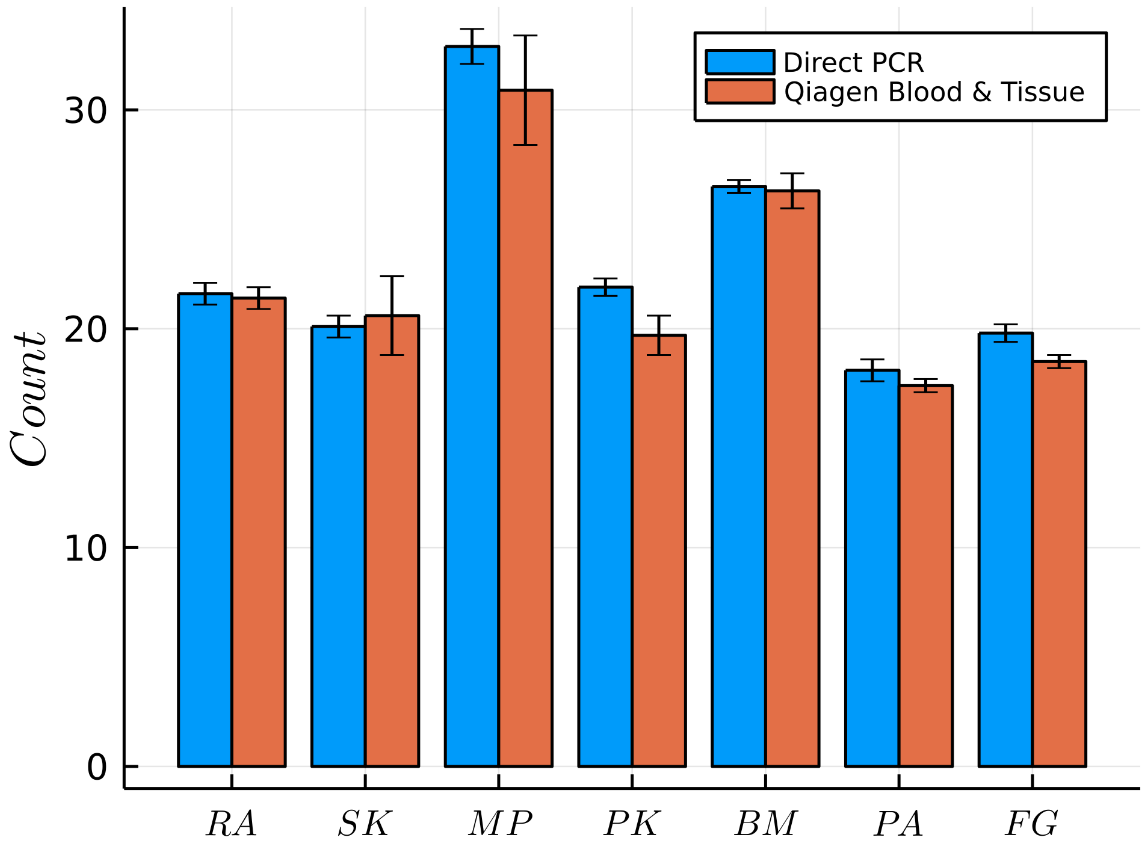


**Figure 12: Validation of Direct PCR** Comparison of the Direct PCR method [30] with standard DNeasy Blood & Tissue kit (Qiagen) counts for each of the soil isolates.

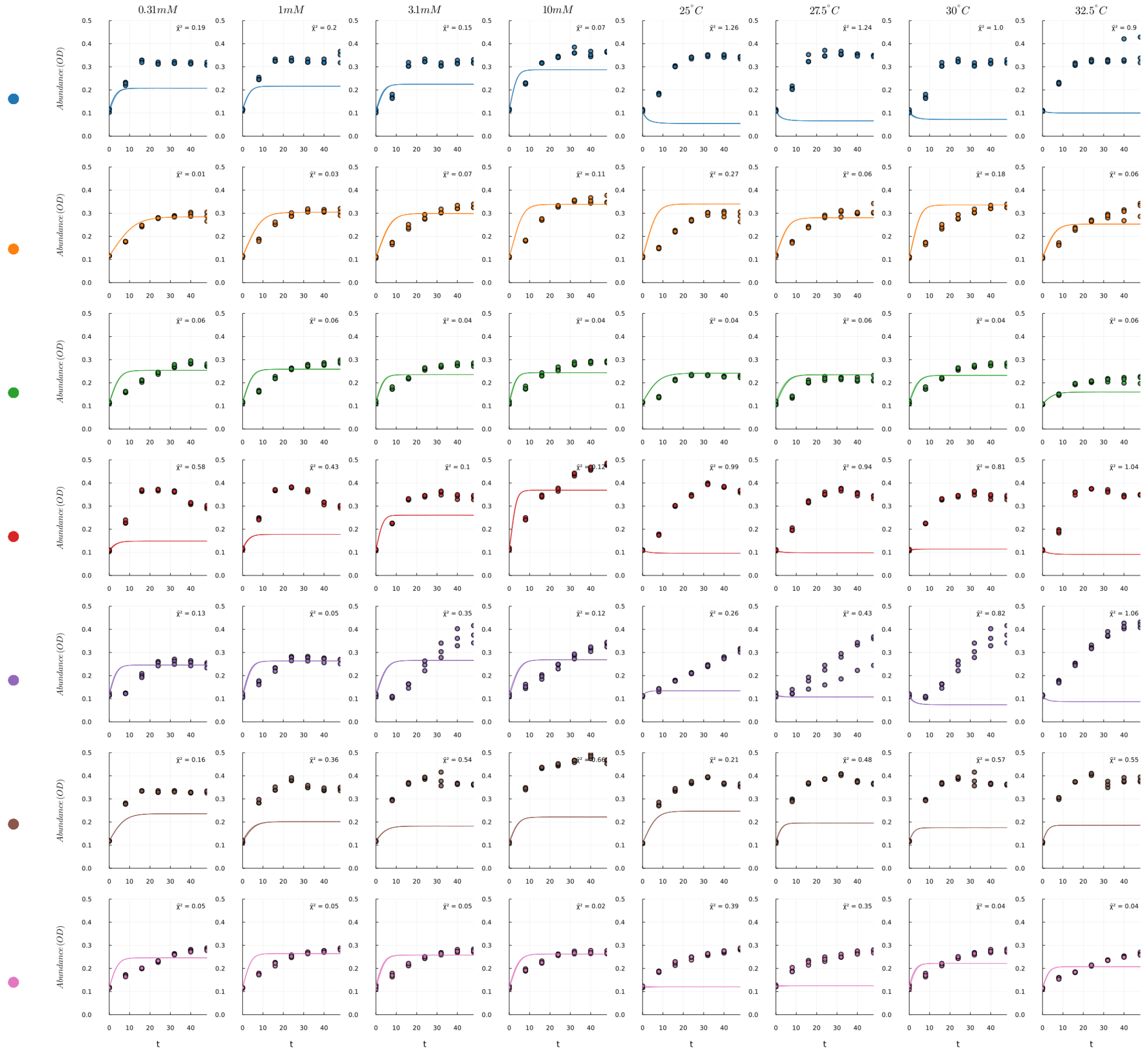


**Figure 13: Model Fit on Isogenic Training Data** The best fit compared with the training data of all isogenic cultures. Real timecourse data are depicted as circles and model prediction is depicted as continuous lines. We note that several of the fits seem poor, however, methods that improved isogenic fits resulted in worse generalization to full community models, perhaps because microbes change their ecological strategy in the presence of other species.

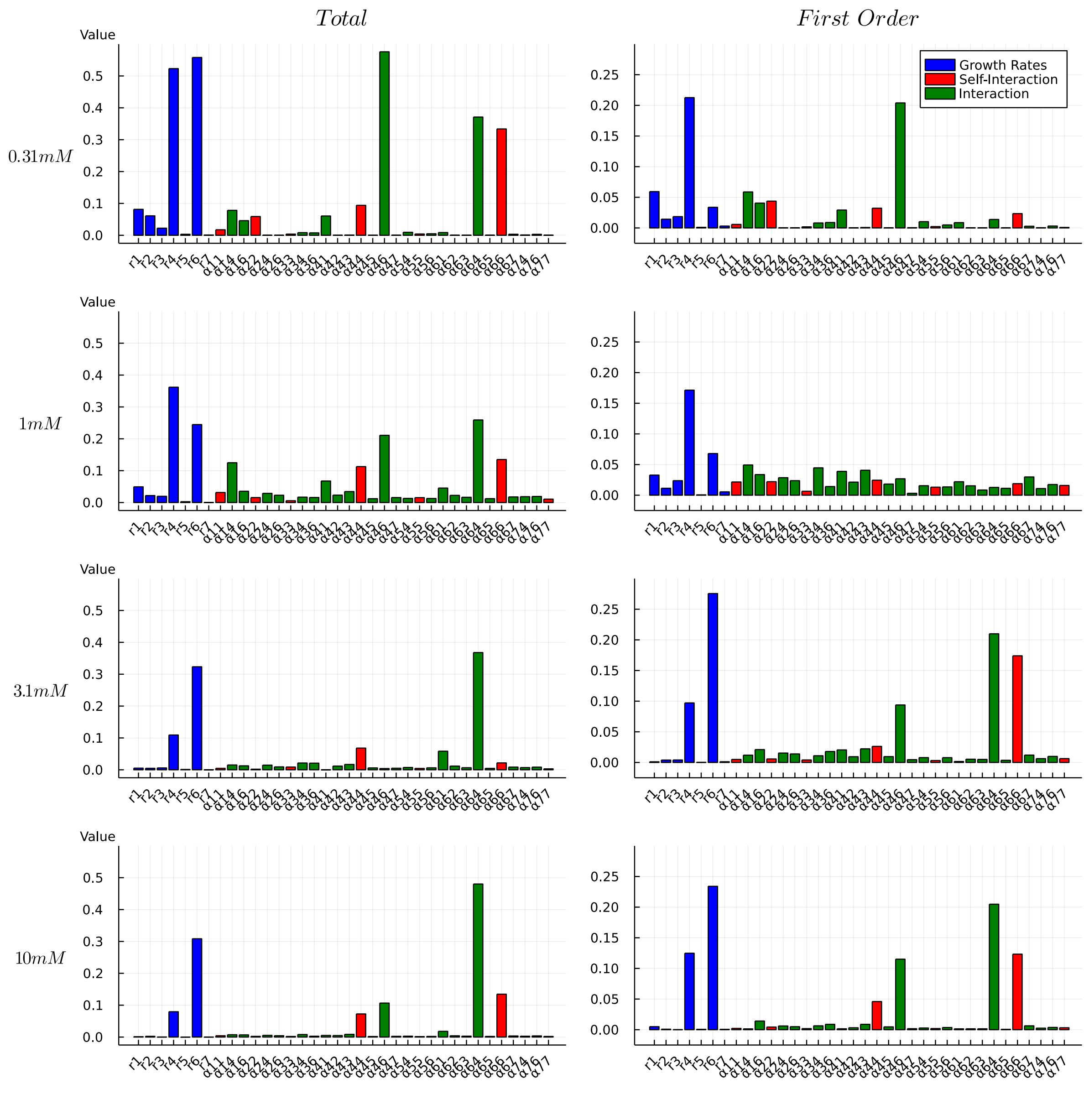


**Figure 14: Sobol Indices of Best Glucose Fits** Total and First order Sobol sensitivity of best fit parameter sets for glucose-fit models with N=1000 samples. Parameter types are classified by color and species numerically labeled corresponding to the order RA, iSK, MP, PK, BM, PA, iFG.

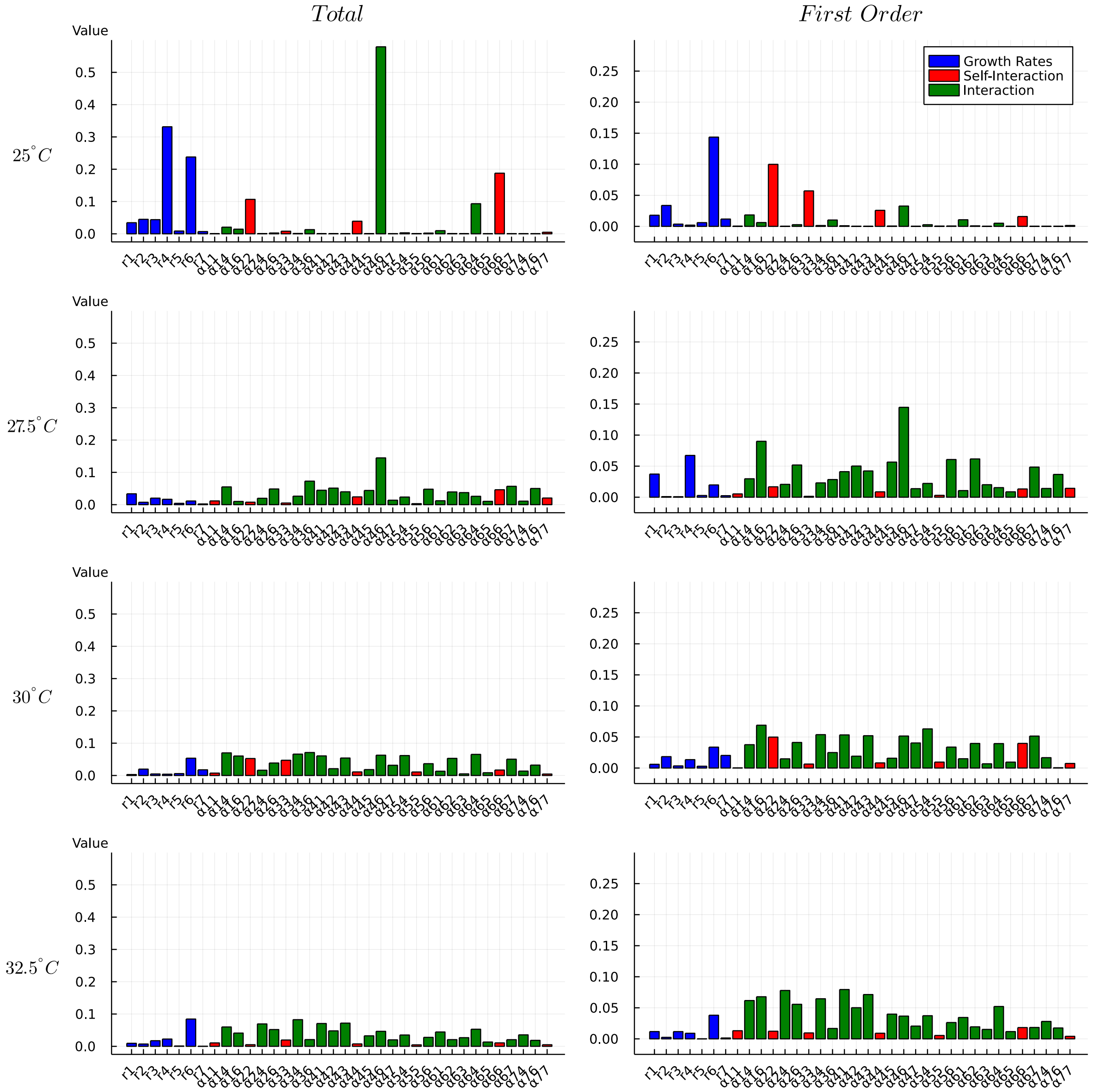


**Figure 15: Sobol Indices of Best Temperature Fits** Total and First order Sobol sensitivity of best fit parameter sets for temperature-fit models with N=1000 samples. Parameter types are classified by color and species numerically labeled corresponding to the order RA, iSK, MP, PK, BM, PA, iFG.

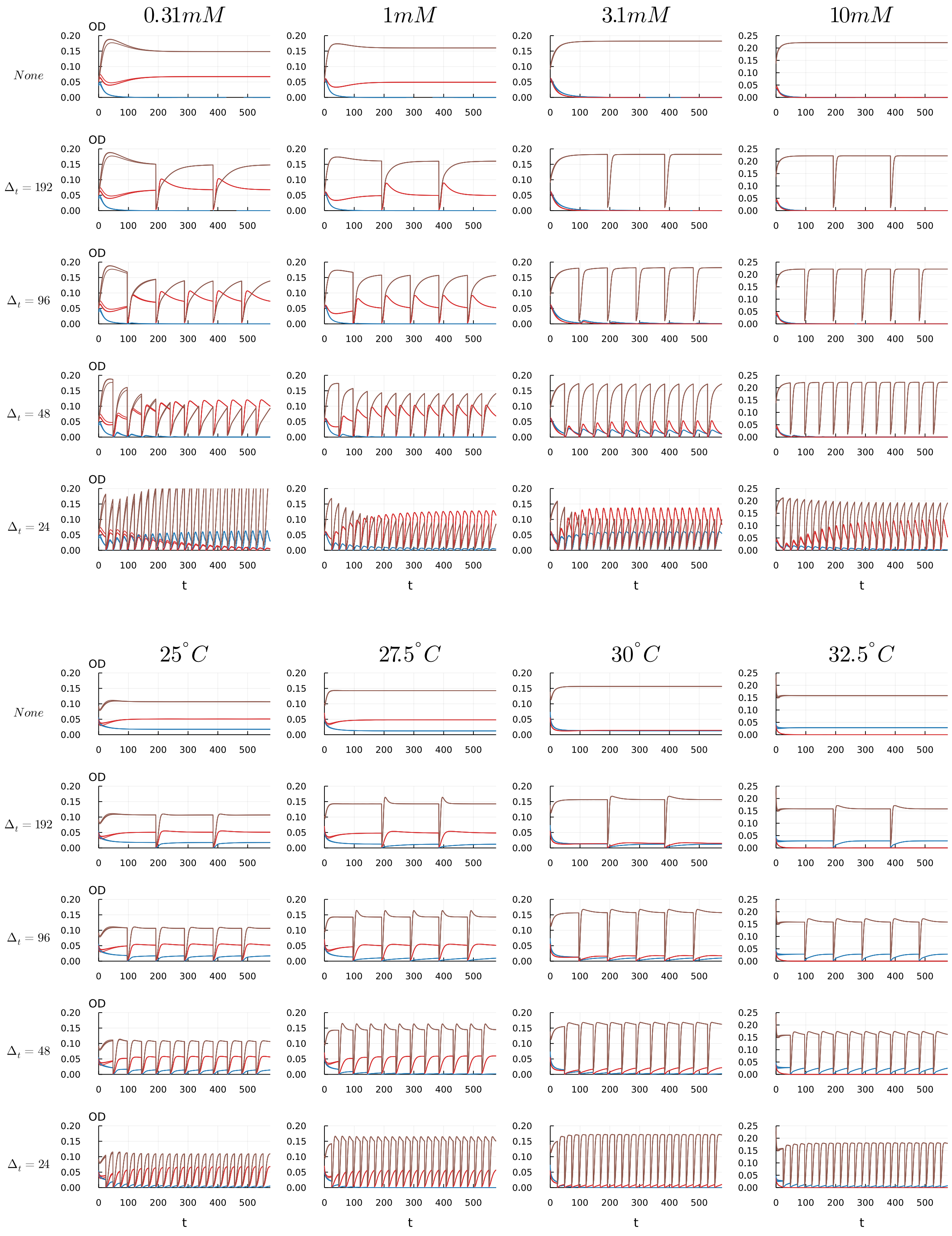


**Figure 16: Effect of Reset Frequency on Community Trajectory** Model-predicted trajectories of the autonomous principle subcommunity, stabilizing to fixed points or cyclic hybrid trajectories for various reset/passaging frequencies (0.05x state reset).

| | $r$ | $A$ | | | $\bar{r}$ | $\bar{\alpha}$ | | $r$ | $A$ | | | $\bar{r}$ | $\bar{\alpha}$ |
|---|---|---|---|---|---|---|---|---|---|---|---|---|---|
| | 0.5 | -2.4 | -3.6 | -2.0 | | | | 0.3 | -5.0 | -2.0 | -0.8 | | |
| 0.31mM | 0.4 | -2.1 | -2.8 | -1.5 | 0.4 | -1.82 | 25C | 0.3 | -4.8 | -3.3 | -0.6 | 0.3 | -2.63 |
| | 0.3 | 0.5 | -1.4 | -1.1 | | | | 0.3 | -3.6 | -2.3 | -1.3 | | |
| | 0.5 | -2.3 | -2.8 | -2.5 | | | | 0.3 | -4.5 | -2.0 | -1.0 | | |
| 1mM | 0.5 | -2.1 | -2.7 | -2.2 | 0.43 | -1.94 | 27.5C | 0.4 | -4.5 | -3.9 | -1.0 | 0.433 | -2.87 |
| | 0.3 | 0.0 | -1.3 | -1.6 | | | | 0.6 | -3.4 | -2.5 | -3.1 | | |
| | 0.5 | -2.2 | -1.4 | -2.9 | | | | 0.3 | -4.4 | -1.8 | -1.5 | | |
| 3.1mM | 0.6 | -2.1 | -2.4 | -3.7 | 0.46 | -2.01 | 30C | 0.4 | -4.4 | -3.7 | -2.0 | 0.433 | -2.93 |
| | 0.3 | -0.8 | -0.8 | -1.8 | | | | 0.6 | -3.3 | -1.8 | -3.5 | | |
| | 0.6 | -2.1 | -1.4 | -2.9 | | | | 0.4 | -4.2 | -1.9 | -1.9 | | |
| 10mM | 0.7 | -2.5 | -2.0 | -3.6 | 0.56 | -2.0 | 32.5C | 0.3 | -4.1 | -3.6 | -1.8 | 0.433 | -2.88 |
| | 0.4 | -0.8 | -0.8 | -1.9 | | | | 0.6 | -3.3 | -1.8 | -3.4 | | |

**Table 3: Parameters of RA,PK,PA from Table 2 and Mean Values**. These parameters govern the principal subcommunity which occupies >99% of total abundance. Mean values of growth rates $\bar{r}$ and interactions $\bar{\alpha}$ are listed for each condition. This subcommunity is used throughout the simulated control experiments.
